# An experimental framework to assess biomolecular condensates in bacteria

**DOI:** 10.1101/2023.03.22.533878

**Authors:** Y Hoang, Christopher A. Azaldegui, Maria Ghalmi, Julie S. Biteen, Anthony G. Vecchiarelli

## Abstract

High-resolution imaging of biomolecular condensates in living cells is essential for correlating their properties to those observed through *in vitro* assays. However, such experiments are limited in bacteria due to resolution limitations. Here we present an experimental framework that probes the formation, reversibility, and dynamics of condensate-forming proteins in *Escherichia coli* as a means to determine the nature of biomolecular condensates in bacteria. We demonstrate that condensates form after passing a threshold concentration, maintain a soluble fraction, dissolve upon shifts in temperature and concentration, and exhibit dynamics consistent with internal rearrangement and exchange between condensed and soluble fractions. We also discovered that an established marker for insoluble protein aggregates, IbpA, has different colocalization patterns with bacterial condensates and aggregates, demonstrating its applicability as a reporter to differentiate the two *in vivo*. Overall, this framework provides a generalizable, accessible, and rigorous set of experiments to probe the nature of biomolecular condensates on the sub-micron scale in bacterial cells.

## Main

Though many cellular compartments and organelles use membranes to encapsulate their contents, biomolecular condensates concentrate proteins and nucleic acids without the use of a membrane^1,2^. Condensates have been widely reported in eukaryotes^2,3^ and more recently in prokaryotic organisms^4,5^, and they have been linked to diverse biological processes important for cell function and physiology^6,7^. Yet, how bacterial condensates maintain their structures, modulate their composition, and regulate internal biochemical reactions is largely unexplored. Our understanding of these questions has improved in eukaryotes^8^, but the small size of bacteria has hindered *in vivo* measurements.

The term condensate is not supposed to prescribe a formation mechanism^2,9^. However, condensate formation has been broadly attributed to phase separation^1,2,7^ because of the liquid-like behavior observed in condensates in living cells and *in vitro* reconstitution. Importantly, condensates that are in a distinct phase from their surrounding environment can be considered as a result of phase separation if they exhibit three key characteristics: a difference in component densities between the two phases, dynamic exchange of components between the two phases, and the formation of a condensate upon reaching a saturation concentration, *c*_sat_^9^. These observations have been made in eukaryotic cells^10–15^ and recently, super-resolution techniques have been implemented to probe sub-diffraction limited bacterial condensates *in vivo*^16–22^. But the general use of these approaches remains limited. The growing number of bacterial systems with reported condensates makes it clear that these membraneless organelles play a key role in bacterial cell biology, making a unifying framework to understand their formation across systems necessary. It is therefore imperative that methods become widely available and flexible to study their formation in bacteria.

Here, we present an experimental framework (Fig. 1) to determine whether a bacterial biomolecular condensate forms through phase separation *in vivo*. We developed and adapted a suite of molecular and cell biology methodologies along with super-resolution imaging techniques in *Escherichia coli* to characterize the formation, solubility, and dynamic exchange of condensates. We examined the protein McdB, a component of the carboxysome positioning system that robustly forms phase-separated droplets *in vitro*^23–25^, alongside well-established control proteins that form condensates^26^ and insoluble aggregates^27,28^. First, we qualitatively probed the liquid-like properties of McdB condensates using overexpression assays. Next, we used tunable expression promoters to quantify the conditions for condensate formation and probed condensate reversibility. Moreover, we measured the dynamic exchange of condensate constituents and inferred the material properties of condensates with single-molecule tracking. Finally, we implemented the heat-shock chaperone IbpA as a molecular sensor that differentiates between condensates and aggregates. Our framework ensures a low barrier to general applicability by users, consolidates an array of adapted and new assays, and complements widely accessible methods with more advanced techniques such as super-resolution imaging and single-molecule tracking to validate method performance.

**Figure 1.**
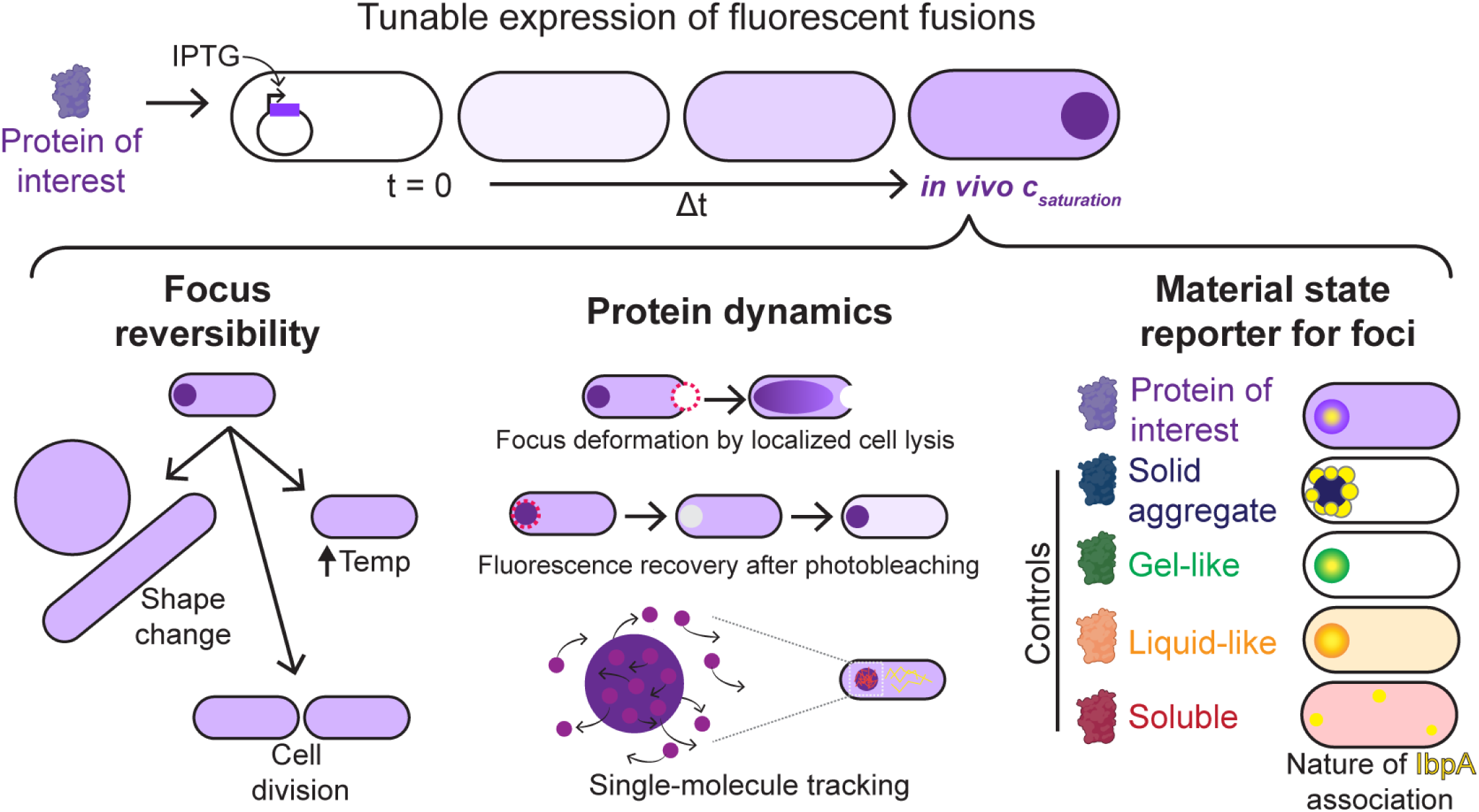
Framework to assess biomolecular condensates in bacteria. Proteins of interest or control proteins are inducibly expressed to form foci (dark purple circle) in *E. coli* (top). The *in vivo* saturation concentration (*c*_saturation_) is quantified. The material state of each focus can be assessed: (*Left)* Condensates dissolve upon cell growth, division, shape change, or temperature shifts, whereas insoluble aggregates remain intact. (*Middle*) Analysis of protein dynamics informs the material properties of the focus. Dashed magenta circles: photobleached areas. (*Right*) The chaperone IbpA (yellow) surrounds insoluble aggregates, but penetrates condensates.

## Results

### Probing the liquid-like properties of foci formed by protein overexpression

Bacterial condensates have been mainly studied *in vitro*, and it is largely unexplored if these proteins form foci with liquid-like properties inside the cell. Overexpression and imaging of fluorescent fusions provide a quick and useful method for gaining insight into the material state of a bacterial focus. The <u>M</u>aintenance of <u>c</u>arboxysome <u>d</u>istribution protein B (McdB) undergoes robust phase separation *in vitro*, but it is unknown whether this activity occurs in the cell^24^. Therefore, we first overexpressed a fully functional mNeonGreen-McdB (mNG-McdB) fusion in *E. coli* BL21 cells and performed time-lapse fluorescence microscopy to qualitatively invesitgate the behaviors of the McdB protein *in vivo*. Focus formation was observed in less than one hour post induction with IPTG at 16 °C (Supplementary Fig. 1a and Supplementary Video 1). Some proximal mNG-McdB foci fused into larger structures on the seconds timescale (Supplementary Fig. 1b). DAPI staining of the nucleoid showed that these larger mNG-McdB foci were nucleoid-excluded (Supplementary Fig. 1c). Strikingly, after three hours, larger mNG-McdB foci localized to the inner membrane near sites of local cell curvature (white arrows in Supplementary Fig. 1a and Supplementary Video 1). We speculate mNG-McdB foci wet to the membrane via nonspecific electrostatic associations and locally occlude cell wall synthesis. The resulting asymmetry in cell wall growth thereby induces cell curvature. Together, we find that mNG-McdB forms small foci upon reaching a threshold cellular concentration, and that these small foci fuse to form large, nucleoid-excluded structures that appear to wet to the inner membrane and cause local changes in cell morphology.

A hallmark of phase-separated condensates is their immediate responsiveness to changes in the cellular environment, such as variations in temperature or cell volume. We first examined the reversibility of mNG-McdB foci via time-lapse microscopy as the stage-top temperature increased from 25 to 37 °C. Despite continued overexpression of mNG-McdB, the foci disappeared within eight minutes, indicative of reversal into a single homogenous phase in the cytoplasm (Supplementary Fig. 1d and Supplementary Video 2). To probe the concentration dependency of mNG-McdB foci formation *in vivo*, we treated the cells with A22 which caused a rod-to-sphere transition and corresponding increase in cell volume. During this transition, mNG-McdB foci dissolved into a homogenous phase in the cytoplasm (Supplementary Fig. 1e, Video 3), suggesting that the cellular levels of mNG-McdB dropped below *c*_sat_.

Because temperature shifts and drug-induced changes to cell volume are gradual and may involve pleiotropic effects, we developed an approach to probe the dynamic properties of mNG-McdB foci upon an instantaneous change to cell volume. In this localized-lysis method, a high-intensity laser was focused on one cell pole to lyse the cell. Localized lysis caused cell contents to unidirectionally rush out of the cell. Strikingly, upon cell lysis, the mNG-McdB focus signal dispersed towards the opposing open cell pole (Supplementary Fig. 1f and Supplementary Video 4). The observed dynamics resemble the observed jetting of P granules under sheer stress^29^. Altogether, these results demonstrate that mNG-McdB foci in *E. coli* BL21 exhibit properties consistent with those of phase-separated condensates.

### Using tunable promoters to probe the formation of biomolecular condensates

In eukaryotic cell biology, the majority of condensate studies perform *in vivo* measurements using ectopic overexpression^30^. However, phase-separating systems are exceedingly sensitive to changes in concentration, and overexpression may introduce significant caveats in the extrapolation that a protein forms condensates when expressed at lower endogenous levels. We set out to find additional metrics other than overexpression to support claims that a bacterial focus is indeed phase-separated.

We turned to a heterologous expression system to observe condensate formation during controlled protein expression. We fused the fluorescent protein mCherry to the N-terminus of McdB(*mCherry-mcdB*) and placed it under the control of an IPTG-inducible promoter on a pTrc99A expression vector (*pTrc99A-mCherry-mcdB*). Along with our protein of interest, we probed a series of well-established control proteins, all fused to mCherry. The protein cI^agg^ is a truncated and mutated version of the Lambda cI repressor that is well-known to form insoluble aggregates in *E. coli*^28^. On the other hand, fluorescent protein fusions to PopTag proteins form condensates via phase separation with tunable material properties that depend on the length of the linker between PopTag and the fusion protein^26^. We engineered two versions of the PopTag fusion: PopTag^SL^ with a short (six-amino acid) GS repeat linker (*pTrc99A-mCherry-L6-PopTag*) and PopTag^LL^ with the native linker of the PopZ protein (78-amino acid) (*pTrc99A-mCherry-L78-PopTag*). Based on the linker lengths, PopTag^LL^-mCherry condensates should be more fluid than PopTag^SL^-mCherry condensates, which we expect to be more viscous or in a gel-like state^26^. Additionally, we probed a solubilized McdB mutant (McdB^sol^), previously shown to be abrogated in its phase separation activity both *in vitro* and *in vivo*^25^. Finally, mCherry alone was used as a control for a completely soluble protein. All mCherry fusion proteins showed a lower degradation level (less than 20%) compared to mCherry alone when expressed with the pTrc99A promoter (Supplementary Fig. 2).

Induced expression of these proteins in *E. coli* cells showed focus formation as a function of increasing protein concentration over time (Fig. 2a-b and Supplemental Fig. 3). After one hour of expression, approximately 60% of mCherry-cI^agg^ and mCherry-PopTag^SL^ cells had a focus, while no foci were observed in mCherry-PopTag^LL^, -McdB, and -McdB^sol^ cells (Fig. 2b). Between one and five hours of expression, the percentage of mCherry-cI^agg^ and mCherry-PopTag^SL^ cells with a focus slightly increased from 60 to 80%, while that of the mCherry-PopTag^LL^ and mCherry-McdB cells significantly increased to 70% and 60%, respectively. The proportion of the mCherry-McdB^sol^ cells with a focus was two-fold lower than that of mCherry-McdB at 5 hours. Strikingly, a notable fraction of the fluorescence signal was localized to the cytoplasm of cells with a mCherry-PopTag^SL^, -PopTag^LL^, -McdB, or -McdB^sol^ focus, but not in the mCherry-cI^agg^ cells (Fig. 2a). The focus and the cytoplasmic fraction resemble the dense phase and dilute phase in a two-phase system.

**Figure 2.**
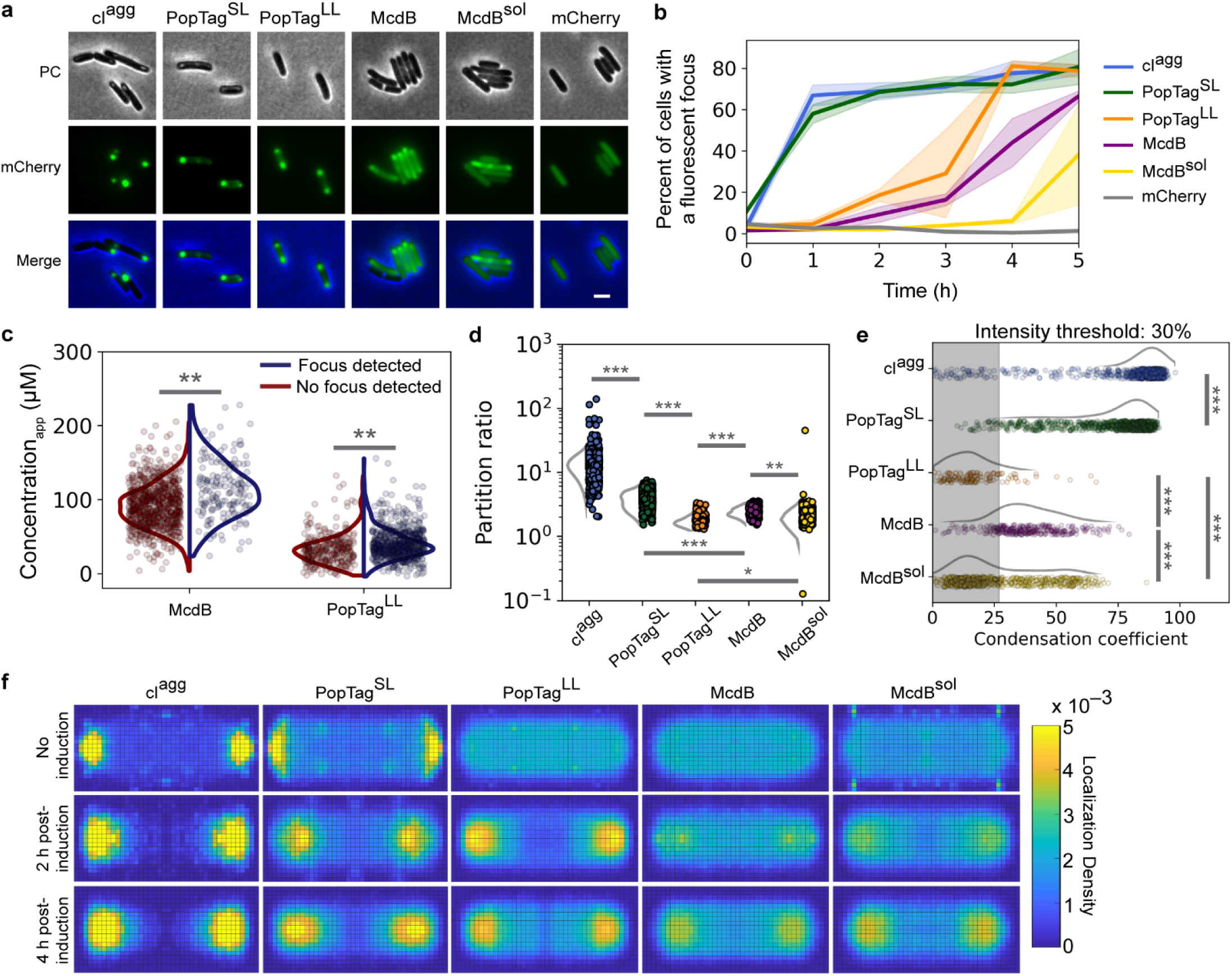
Proteins that phase separate *in vitro* maintain a soluble cytoplasmic fraction and form foci above a quantifiable *in vivo c*_sat_. **a.** Representative images of mCherry fusion protein foci in *E. coli* at 4 h post-induction. Phase contrast (PC) (gray), mCherry channel (green), and merged images are shown. Images are representative of three biological replicates. Scale bar: 2 µm. **b.** Percent of cells with a focus over induction time. Colored solid lines and shading represent the average and standard deviation, respectively, over three biological replicates. *N* > 100 cells for each protein at each timepoint. **c.** Quantification of an apparent cellular *c*_sat_. Cells are classified by the presence (blue) or absence (red) of a focus—above or below the apparent concentration *c*_sat_app_ respectively (834 cells without an McdB focus, 170 cells with an McdB focus, 311 cells with a PopTag^LL^ focus, 681 cells without a PopTag^LL^ focus). Data points correspond to individual cells. The curves next to the scatter plots were obtained via kernel density estimation. Statistical analysis was done on a set of 100 randomly selected cells for each classification. **d.** Partition ratios for specified mCherry fusions with detected foci (cI^agg^, *N* = 1013 cells; PopTag^SL^, *N* = 893 cells; PopTag^LL^, *N* = 151 cells; McdB, *N* = 284 cells; McdB^sol^, *N* = 717 cells; mCherry, *N* = 53 cells). **e.** Single-cell condensation coefficients for specified mCherry fusions with detected foci. Data points correspond to individual cells. The curves next to the scatter plots were obtained via kernel density estimation. The shaded region represents the measurement range for cells expressing a uniform mCherry signal. *N* values are same as in (d). **f.** Normalized single-molecule localization 2D histograms. Single-molecule localizations of specified PAmCherry fusion proteins were collected, projected, and binned onto a normalized cell shape to generate the heat map. *N* denotes the number of cells analyzed for each strain over three biological replicates. *p < 0.05, \*\**p* < 0.01, and ***p < 0.001 by Welch’s *t* test.

Next, we used quantitative fluorescence microscopy to determine the apparent *in vivo* saturation concentration (*c*_sat_app_) of our protein of interest, McdB. We first determined the intensity of single mCherry molecules that were spatially isolated prior to photobleaching of fluorescence in the cell. These single-molecule localization images were quantified to measure the number of photons detected per molecule per imaging frame (Supplementary Fig. 4). Next, we calculated the cellular concentration of McdB by integrating the total cellular fluorescence intensity per imaging frame and dividing this value by the mCherry single-molecule intensity and the cellular volume. Cells were then classified by the presence or absence of a focus. After a 4-hour induction, cells expressing mCherry-McdB without a detected focus had an average cellular concentration of 92 ± 29 µM, while the concentration in cells with a focus was 113 ± 37 µM (Fig. 2c). This observation of condensates in cells with a slightly higher total protein concentration is consistent with McdB undergoing a nearly immediate and concentration-dependent transition to form a focus. We performed the same analysis for cells expressing mCherry-PopTag^LL^. The average cellular concentration of this fluid-condensate control was 32 ± 20 µM in cells without a focus while cells with a detected focus had an average concentration of 40 ± 20 µM (Fig. 2c). PopTag^SL^ formed condensates prior to IPTG induction, which prevented similar analyses. Together, the results provide a *c*_sat_app_ at which condensates form; for mCherry-PopTag^LL^ and mCherry-McdB, we estimate the *c*_sat_app_ to be between 32 – 40 µM and 92 – 113 µM, respectively.

### Condensates coexist with a soluble phase

A hallmark of phase separation is the existence of a soluble fraction in the cytoplasm. To quantify the ratio of mCherry protein fusion concentration in the focus (dense phase) to the concentration in the cytoplasmic fraction (dilute phase), the partitioning of each fusion was measured (Fig. 2d). The largest partitioning was measured for the aggregator control, mCherry-cI^agg^, in which there was no detectable fluorescent signal in the cytoplasm. The mCherry-PopTag^SL^ condensate control partitioned to a greater extent than the more fluid condensate control mCherry-PopTag^LL^. The partitioning of our protein of interest, mCherry-McdB, was intermediate relative to the PopTag controls. We also analyzed condensation in cells by normalizing all pixel intensities in hundreds of cells per protein and calculating the fraction of pixels below a normalized intensity threshold of 30% (Supplementary Fig. 5)^16,31^. The relative condensation coefficients acquired from this analysis (Fig. 2e) were consistent with the partition ratios (see Fig. 2d).

To further inspect the partitioning of proteins to polar foci, we titrated photoactivatable (PA) mCherry fusions of each protein and performed single-molecule localization super-resolution microscopy to generate normalized localization density heat maps (Fig. 2f). Consistent with our previous results, PAmCherry-cI^agg^ had minimal localization density in the cytoplasm under all conditions, which suggests nearly all protein is recruited to the polar aggregates. On the other hand, the polar density of PAmCherry-PopTag^SL^ increased with increasing protein concentration, but these cells also maintained a protein fraction in the cytoplasm. Strikingly, PAmCherry-PopTag^LL^ and -McdB displayed a transition between no induction and two hours post induction, indicative of focus formation. Intriguingly, PAmCherry-McdB^sol^ also formed high-density regions at the poles, consistent with bulk fluorescence measurements (see Fig. 2a). To abrogate phase separation in this McdB^sol^ mutant, the net negative charge of its IDR was increased^25^; therefore, we speculate that the localization pattern of the fluidized PAmCherry-McdB condensate is due to nucleoid exclusion by repulsive electrostatic interactions. Consistently, we found that mCherry-McdB^sol^ foci expanded and encroached throughout the cytoplasm as the nucleoid was compacted via drug treatment (Supplementary Fig. 6 and Supplementary Video 5). Collectively, our results demonstrate the use of a tunable expression system that shows the concentration-dependent formation of condensates that exhibit a two-phase behavior, indicative of phase separation. Phase separation theory predicts that while many small condensates form at the initiation of phase separation, this number decreases, and the sizes of those that persist increase via coalescence^32,33^.

Our findings here are consistent with the expected final ground state being a single large condensate that coexists with a dilute phase. Some cells had foci at opposing poles. However, this can be attributed to the steric effects of nucleoid exclusion that prevent condensates from physically interacting and coalescing.

### Bacterial cell growth and division probe the reversibility of condensates

Condensates should dissolve if cellular levels of the protein drop below *c*_sat_ ^7^. To probe condensate reversibility in bacteria, we implemented two approaches that lower the protein concentration in the system following focus formation. We expected that driving the protein concentration below the protein *c*_sat_ would dissolve the condensates and thus demonstrate solubility as a driving force for condensate formation. First, each protein was expressed via IPTG induction, and once foci formed, expression was stopped to maintain a constant cellular protein level. Cells were then allowed to grow and divide, and in doing so, dilute the concentration of the mCherry fusion proteins. Noticeably, mCherry-cI^agg^ foci remained at the pole from which they formed even as cells grew and divided over 15 hours (∼9 generations), resulting in an average focus lifespan of 14.3 ± 2.2 hours (Fig. 3a-c and Supplementary Video 6). This result indicates that there is no concentration dependence in the formation and maintenance of mCherry-cI^agg^ foci, and further supports the previously reported insolubility of cI^agg^ foci^28^.

**Figure 3.**
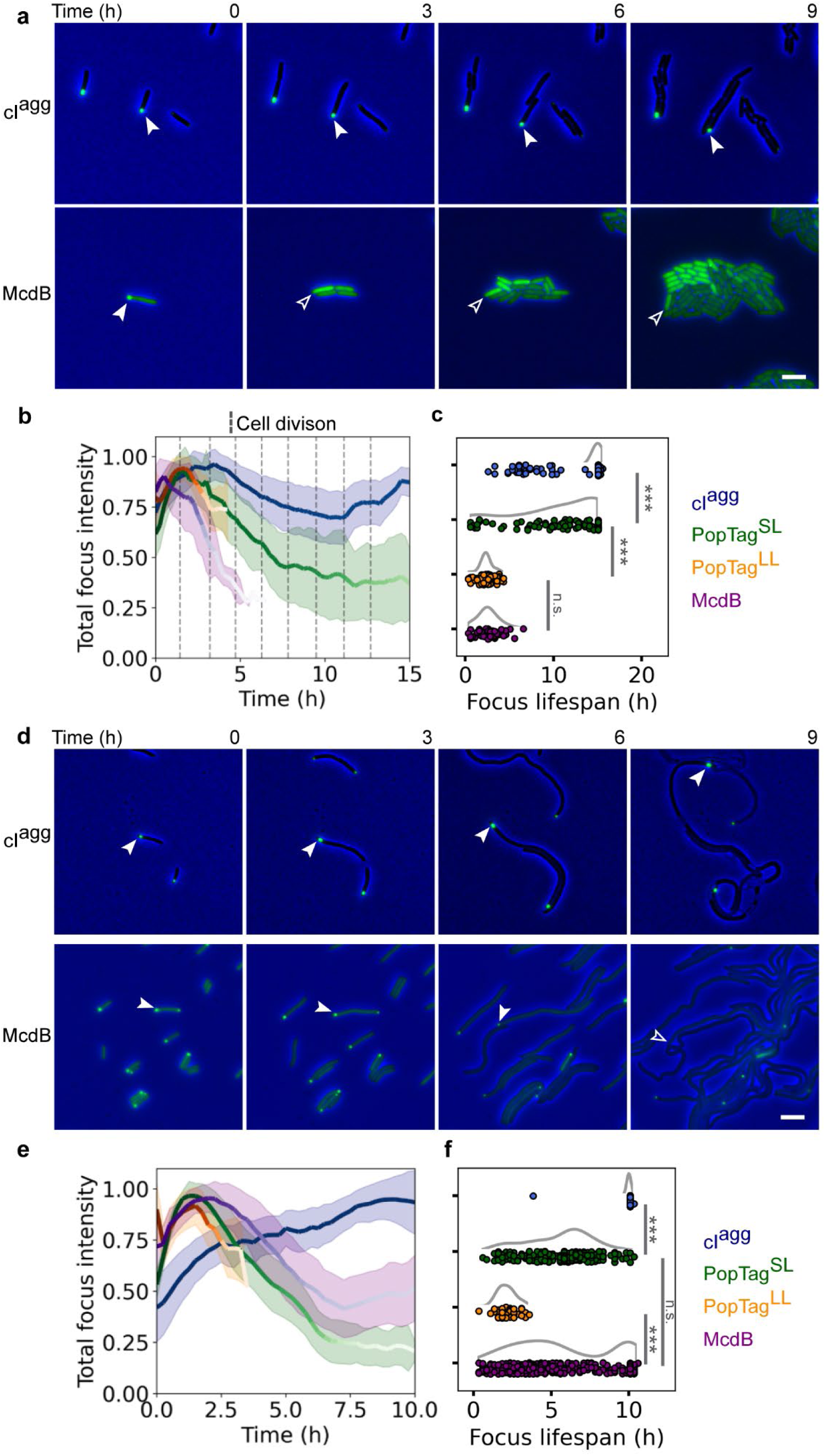
Bacterial cell growth and division dissolve condensates by dropping the cellular concentration below the apparent *c*_sat_. **a.** Generational-dilution dissolves condensates. The phase contrast channel (blue) and the mCherry channel (green) were merged and shown for indicated time points. White arrows demarcate the cellular location of the same focus over time. Blank arrows demarcate the same cellular position now absent of a focus. Images are representative of four biological replicates. Scale bar: 2 µm. **b**. Quantification of focus intensity through cell generations. Colored solid lines and shading are the average and standard deviation, respectively, of the normalized total focus intensity for each strain indicated. The solid line color gradient over time indicates a decrease in the total number of foci detected at that indicated time point due to dissolution. cI^agg^: *n* = 427 foci; PopTag^SL^: *n* = 98 foci; PopTag^LL^: *n* = 81 foci; McdB: *n* = 48 foci. Dashed vertical lines indicate cell division events. **c.** Lifespan of protein foci during cell division. Lifespan of individual foci was determined by the particle trajectory length (see Methods). Data points correspond to individual foci. The curves next to the scatter plots are obtained via kernel density estimation. \*\*\**p* < 0.001 by Welch’s *t* test; n.s. indicates no statistically significant difference between the samples. **d.** Cell elongation dissolves phase-separated condensates. As in (a), except cells were treated with 10 µg/ml cephalexin to block cell division. Scale bar: 2 µm. **e.** Quantification of focus intensity during cell elongation. Color scale as in (b). cI^agg^, *n* = 210 foci; PopTag^SL^, *n* = 239 foci; PopTag^LL^, *n* = 44 foci; McdB, *n* = 464 foci. **f.** Lifespan of protein foci during cell elongation. Same as in (c).

In contrast, mCherry-PopTag^SL^, mCherry-PopTag^LL^, and mCherry-McdB foci all exhibited concentration-dependent dissolution. mCherry-PopTag^SL^ foci had an average lifespan of 11.0 ± 4.4 hours (Fig. 3b-c, Supplementary Fig. 7a), and the total focus intensity dropped approximately two-fold over this time period, suggesting a decrease in focus size. The generational dilution effect was more immediate with the mCherry-PopTag^LL^ (Supplementary Fig. 7a), which exhibited a rapid decrease in total focus intensity and full dissolution within one to three cell divisions and an average focus lifespan of 2.3 ± 0.8 hours. When compared to these controls, mCherry-McdB focus dissolution mirrored that of the fluid mCherry-PopTag^LL^ condensate, with an average focus lifespan of 2.6 ± 1.3 hours or within one to three cell divisions. Indeed, many mCherry-McdB foci dissolved following a single cell division event (Supplementary Video 6). Interestingly, we observed some mCherry-McdB and mCherry-PopTag^SL^ foci reforming immediately after cell division (Supplementary Fig. 8). This observation is consistent with the low *c*_sat_ of mCherry-McdB and mCherry-PopTag^SL^. As the cell divides, the volume decreases, leading to an increase in the concentration back above *c*_sat_ in the daughter cell, which ultimately results in the reformation of a focus.

We also probed condensate solubility by diluting protein concentration via an increase in cell length without further protein expression. *E. coli* cells were prepared as described above, but with the inclusion of the cell division inhibitor cephalexin (10 µg/ml). We expected similar trends for dilution via cell elongation as in cell division, and indeed mCherry-cI^agg^ foci were observed to persist throughout the duration of the experiment (10 hours), with an average lifespan of 10.0 ± 0.4 hours (Fig. 3d-f and Supplementary Video 7). The total focus fluorescence of mCherry-PopTag^SL^ foci, on the other hand, decreased significantly over time and completely dissolved with an average lifespan of 5.5 ± 0.4 hours (Fig. 3e-f, Supplementary Fig. 7b). The more fluid PopTag control, mCherry-PopTag^LL^, readily dissolved with an average lifespan of 2.0 ± 0.6 hours.

Strikingly, the average lifespan of mCherry-McdB foci was 5.6 ± 3.1 hours, which was roughly twice as long as that found in the cell division experiments above. This increase in average lifetime was influenced by the persistence of some foci over the duration of imaging, which resulted in a bimodal distribution of foci lifespan for mCherry-McdB (Fig. 3f). The reason for this persistence of some foci is unknown. But we speculate that the altered cytoplasmic conditions (e.g. effective volume and crowding) of multi-nucleate elongated cells, compared to cells undergoing vegetative growth, play a role in the observed differences in focus dissolution.

Together, using our assays of protein dilution via cell division or cell elongation, we observed decreases in focus size toward its eventual dissolution, and a corresponding turnover of protein into the cytoplasmic phase. The findings are consistent with the reversal to a one-phase system once the concentration has decreased below *c*_sat_. These characteristics further support condensate formation by the PopTag fusions via phase separation and suggest a similar formation process for McdB condensates.

### Probing the dynamic rearrangement, confinement, and exchange of biomolecular condensates

To probe the dynamic rearrangement of molecules within a focus and their exchange with the surrounding cytoplasm, we first implemented fluorescence recovery after photobleaching (FRAP) on the mCherry fusion foci. The aggregator control, mCherry-cI^agg^, exhibited no fluorescence recovery (Fig. 4a-b), supporting the static nature of the proteins within these aggregates and the absence of protein exchange with the cytoplasm. The PopTag fusions partially recovered to an extent consistent with their respective fluidity levels: mCherry-PopTag^SL^ and -PopTag^LL^ recovered to approximately 10% and 40%, respectively, of the initial fluorescence intensity (Fig. 4b). Recovery of mCherry-McdB foci was once again similar to that of the fluid mCherry-PopTag^LL^ condensate. Taken together, the data suggest that McdB exhibits dynamics similar to that of the fluid PopTag^LL^, while the minimal recovery of PopTag^SL^ is consistent with its predicted gel-like state. Given the limited number of pixels that make up the photobleached foci, we note that the fluorescence recovery contributions from internal rearrangements of molecules within a focus cannot be distinguished from exchange of molecules with the surrounding milieu^30^. While FRAP is commonly used to determine if a compartment is liquid-like, this method is solely a measure of exchange dynamics, so FRAP alone is not a reliable “gold standard” measure of the material state of a focus.

**Figure 4.**
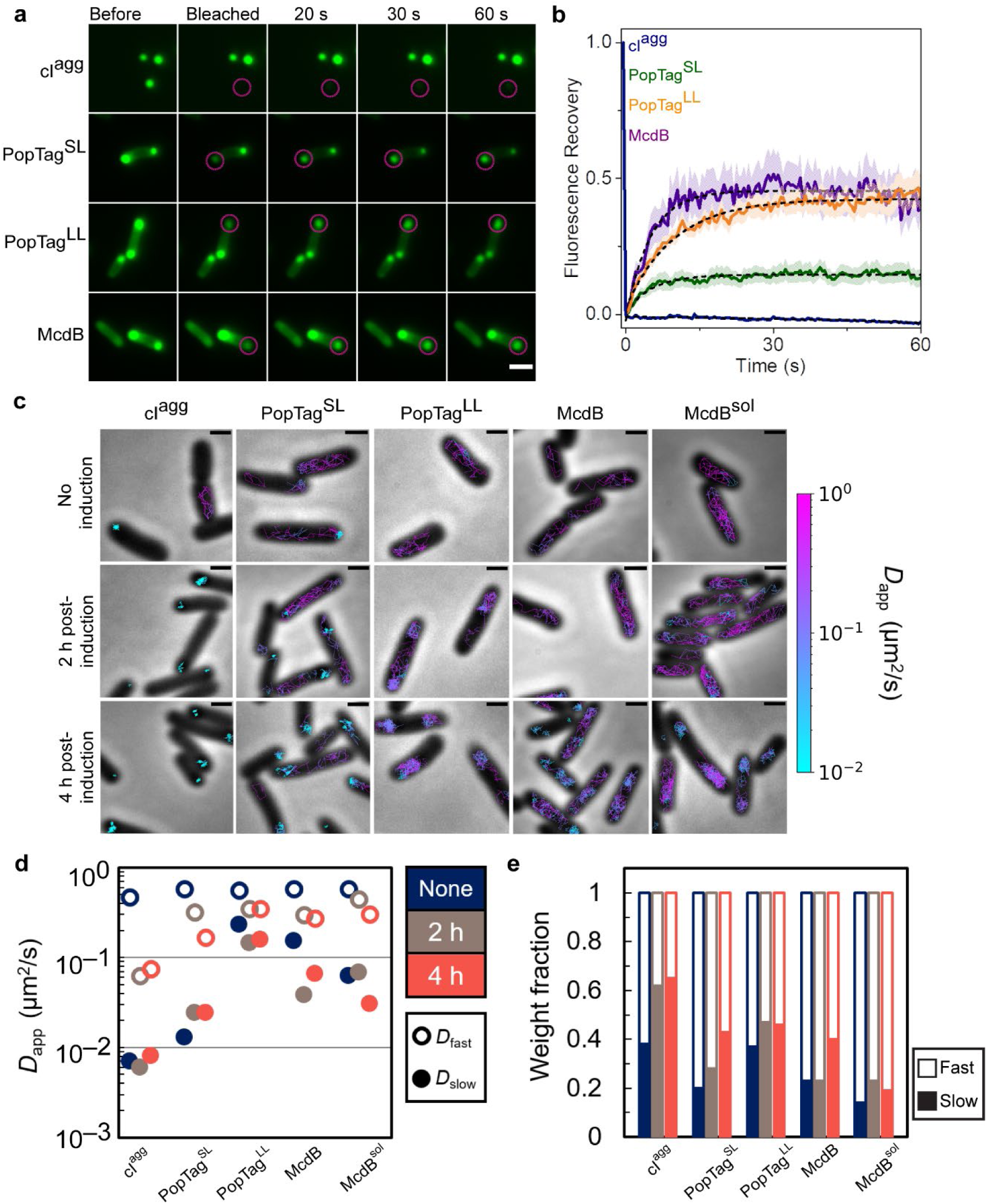
FRAP in concert with single-particle tracking informs the material state of foci in bacterial cells. **a.** Fluorescence recovery of mCherry-fusion proteins after photobleaching. Magenta circles indicate FRAP region. Scale bar: 2 µm. **b.** Quantification of fluorescence recovery of mCherry fusion proteins. Shading represents the standard error of the mean. cI^agg^: *n* = 14 foci; PopTag^SL^: *n* = 12 foci; PopTag^LL^: *n* = 14 foci; McdB: *n* = 10 foci. **c.** Representative single-molecule trajectories. Overlay of tracks, color-coded according to the track apparent diffusion coefficient, obtained from representative *E. coli* cells expressing PAmCherry fusion proteins. Scale bars: 1 µm. **d.** Diffusion coefficients of PAmCherry fusion proteins. The diffusion coefficients of the indicated PAmCherry fusions were obtained by fitting the histograms of the log diffusion coefficients of single tracks to a two-component Gaussian mixture model assuming that the fusion proteins are composed of a slow (closed circles) and fast (open circles) fraction. **e.** Weight fractions of mobility states of PAmCherry fusion proteins. Two-component Gaussian mixture fitting results show an increase in slow mobility fraction (solid bars) and decrease in fast mobility fraction (empty bars) as protein concentration increases.

Therefore, to further investigate the dynamics of our set of proteins both in the focus and in the cytoplasm, we implemented single-molecule localization microscopy and tracked the movement of individual molecules. We then determined the apparent diffusion coefficients (*D*_app_) of the PAmCherry fusions to the screened proteins under different induction conditions by measuring the mean square displacement (MSD) of individual trajectories as a function of time lag, *τ*. Prior to induction, low-level leaky expression of all fusions displayed a fast diffusive state (*D*_app, fast_) (Supplementary Fig. 9). Trajectory mapping back onto the cell showed the *D*_app, fast_ population corresponds to free diffusion in the cytoplasm (Fig. 4c). PAmCherry-cI^agg^ and -PopTag^SL^ also displayed a small population of nearly static molecules (*D*_app, slow_) (Supplementary Fig. 7), with trajectories that mapped to the cell poles (Fig. 4c). Upon induction, the near-static fraction of molecules increased for PAmCherry-cI^agg^ and -PopTag^SL^, and became the dominant state for PAmCherry-cI^agg^ (Supplementary Fig. 9). A diffuse state, slower than free diffusion, also emerged for PAmCherry-McdB, which mapped to the cell poles (Fig. 4c). The vast majority of PAmCherry-McdB^sol^ molecules, on the other hand, remained in the fast diffusive state but remained nucleoid excluded (Fig. 4c); consistent with our wide-field microscopy of this fluidized McdB mutant (Supplementary Fig. S5). PAmCherry-PopTag^LL^ also exhibited nucleoid exclusion (Fig. 4c), despite remaining highly mobile. As stated earlier, we speculate that this behavior is caused by charge repulsion with the nucleoid.

By comparing the average mobility of *D*_app, slow_ populations across all fusions, we find that PAmCherry-cI^agg^ molecules within a focus were essentially static (Fig. 4d), consistent with being an aggregate. PAmCherry-PopTag^SL^ molecules in foci displayed an intermediate mobility consistent with a gel or highly-viscous fluid state. All PAmCherry-PopTag^LL^ molecules, on the other hand, essentially displayed a monomodal diffusive distribution across all induction conditions, consistent with this version of the PopTag being highly fluid.

Finally, PAmCherry-McdB displayed a *D*_app, slow_ state only slightly less mobile than the *D*_app, fast_ population, consistent with a liquid-like condensate. Although our fluidized mutant of McdB, PAmCherry-McdB^sol^ displayed a *D*_app, slow_ population similar to that of PAmCherry-McdB, *D*_app, slow_ was a minor fraction of the total population (Fig. 4e and Supplementary Figure S7). Collectively, the data parse the dynamic exchange of molecules in foci and the surrounding cytoplasm, and delineate the spectrum of possible mobilities within a focus, thus allowing for inference of its material state.

### IbpA as a reporter to differentiate between condensates and aggregates

The heat shock chaperone, IbpA, has been shown to colocalize with insoluble aggregates in *E. coli*^28^. We therefore initially thought that IbpA may work as an *in vivo* reporter that discriminates between aggregates and condensates, colocalizing with the aggregator control and not recognizing PopTag or McdB condensates. We expressed the mCherry fusion proteins under focus-forming conditions in an *E. coli* strain that expressed a chromosomal fluorescent reporter of IbpA (IbpA-msfGFP) and then observed colocalization patterns of the mCherry fusion proteins with IbpA. Unexpectedly, the IbpA foci strongly colocalized with all proteins surveyed (Supplementary Fig. 10a), showing that IbpA does not specifically localize to misfolded protein aggregates in the cell as ascribed by current models^28^. Upon further investigation, we observed that the localization pattern of IbpA association with the aggregate control versus the condensates was quantifiably different. We quantified and compared the diameters of the IbpA signal projections relative to each mCherry fusion (Supplementary Fig. 10b) and found that the IbpA signal was spread over a larger area than the mCherry-cI^agg^ foci. We also found that in many instances IbpA appeared to coat, as opposed to penetrate, the mCherry-cI^agg^ foci (Supplementary Fig. 10a).

We extracted from the wide-field fluorescence data an intensity profile line plot of foci in both color channels. When normalized to the maximum intensity of the mCherry signal, we observed significant differences in the relative amounts of IbpA penetrating each of the mCherry fusion foci (Fig. 5a). On average, the max intensity of IbpA was 20 ± 9 % of that of mCherry-cI^agg^ foci, while it was 58 ± 33, 188 ± 131, and 33 ± 35 % for mCherry-PopTag^SL^, -PopTag^LL^, and -McdB foci, respectively. The limited amount of IbpA relative to cI^agg^ suggested to us that while IbpA can sense the aggregate, it cannot penetrate it well. In contrast, the higher amounts of IbpA in all other foci pointed to its ability to penetrate more fluid assemblies. This result demonstrates varying degrees of IbpA penetration within the corresponding mCherry fusion foci which correlate well with their hypothesized material state and suggest a different mode of colocalization of IbpA with biomolecular condensates versus aggregates.

**Figure 5.**
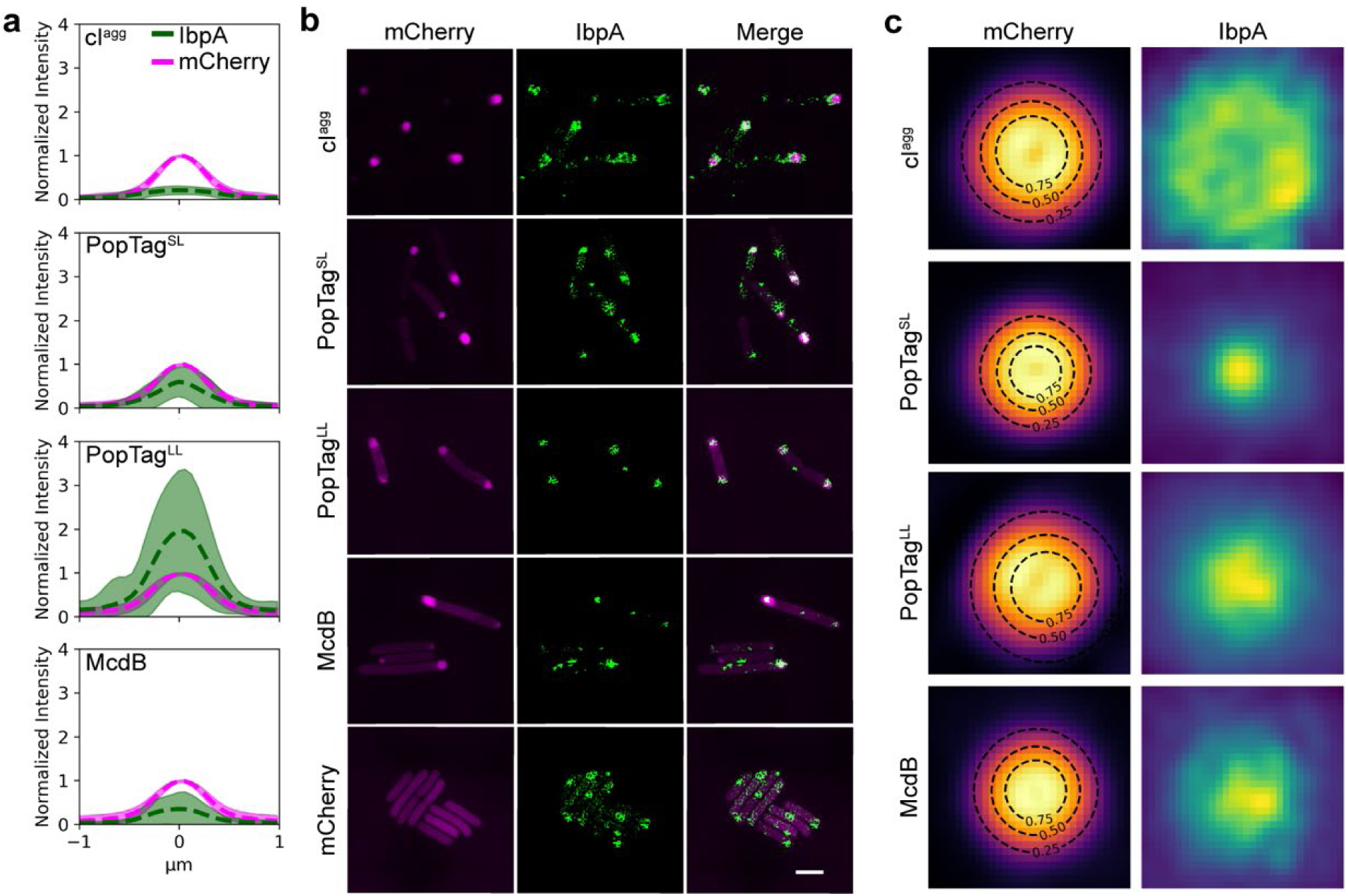
Nature of IbpA association with mCherry fusion foci by 2D SIM. **a.** Fluorescence intensity profile lines of mCherry fusion foci (magenta) and IbpA-msfGFP foci (green). Dashed lines and shading are the mean and standard deviation, respectively, of fluorescence intensities normalized to mCherry signal. cI^agg^: *n* = 460 foci; PopTag^SL^: *n* = 685 foci; PopTag^LL^: *n* = 874; McdB: *n* = 332 foci. **b**. 2D SIM images of mCherry fusion proteins foci (magenta) and IbpA foci (green) are shown. Images are representative of three biological replicates. Scale bar: 2 µm. **c.** Fluorescence projections of mCherry fusion foci and local IbpA-msfGFP signal. Black dashed lines indicate the contour at 0.25, 0.50, and 0.75 of the 2D Gaussian fit amplitude surrounding the mCherry fusion focus. cI^agg^: *n* = 84 foci; PopTag^SL^: *n* = 506 foci; PopTag^LL^: *n* = 457 foci; McdB: *n* = 217 foci.

To better resolve the colocalization patterns between IbpA and the mCherry fusion foci, we imaged cells by 2D structured illumination microscopy (SIM) under the same conditions as described earlier. With the increased resolution in both channels, the localization patterns of IbpA relative to the mCherry fusion foci were more easily observed (Fig. 5b). Using a projection of all detected foci for each mCherry fusion superimposed onto one another, we observed that IbpA forms a rosette pattern around the mCherry-cI^agg^ foci, a punctate focus within mCherry-PopTag^SL^ foci, and an amorphous focus pattern with mCherry-PopTag^LL^ and mCherry-McdB (Fig. 5c). This result demonstrates that IbpA exhibits a different colocalization pattern with biomolecular condensates compared to aggregates and better penetrates more fluid condensates. These patterns serve as a proof of concept that IbpA is a reporter capable of differentiating between these macromolecular assemblies *in vivo*.

## Discussion

In this study, we developed an experimental framework to assess the material state of fluorescent foci in bacteria; specifically, whether a focus can be described as a phase-separated condensate. Inducer-controlled protein expression combined with quantitative fluorescence microscopy was shown to have general applicability in identifying the *in vivo c*_sat_ for condensate formation. Decreasing cellular protein levels back below *c*_sat_, via changes in cell shape or by generational dilution, can then probe the reversibility of condensate formation. We further show that protein dynamics within both the cytoplasm and the focus can infer its material state. Finally, we identified the heat-shock chaperone, IbpA as a molecular sensor that surrounds solid aggregates but penetrates condensates. We demonstrated the versatility of these approaches by using foci-forming control proteins that span the spectrum of material states from a solid aggregate to a highly fluid condensate. When compared to these control proteins, we find that our protein of interest, McdB, robustly phase-separates into liquid-like condensates *in vivo*. As shown with the fluidized mutant of McdB, this framework can also be combined with mutagenesis studies to determine the regions and residues of a protein governing its material state and phase separation behavior in a bacterial cell.

Our framework overcomes current limitations in identifying the material state of a fluorescent focus and can be used to assess the phase separation activity of expressed recombinant proteins and bacterial inclusion bodies (IBs). IBs are mesoscale protein aggregates, once strictly proposed as being composed of nonfunctional and misfolded proteins^34^. Phase separation has only recently been considered in the assembly and organization of IBs^35^. For example, several condensates have been shown to mature into gels and solid amyloids^1^. However, direct determination of whether a bacterial IB is a liquid, gel, solid, or a mixture of these states remains to be demonstrated.

Inclusion body <u>b</u>inding <u>p</u>rotein A (IbpA) of *E.coli* belongs to the conserved family of ATP-independent small heat shock proteins, well-established in binding protein aggregates and driving them towards reactivation-prone assemblies^36–38^. As such, we presumed IbpA would serve as a molecular sensor that would selectively associate with protein aggregates, but not condensates. Instead, we found that IbpA surrounded protein aggregates and penetrated condensates. Moreover, the degree to which IbpA colocalized with condensates strongly correlated with increasing fluidity. Consistent with our findings, the Drummond group has recently shown that condensates are dispersed by chaperones far more rapidly than misfolded aggregates^39^. These findings warrant a reevaluation of the function of chaperone systems governing protein homeostasis and demonstrate the utility of IbpA, and potentially other chaperones, as molecular sensors for the material state of fluorescent foci in bacteria.

The framework was built by contrasting the principles of aggregation versus phase separation behaviors of proteins. While not an exhaustive list, the approaches used here are highly accessible and probe several aspects of condensate assembly, maintenance, and dissolution. The framework is not without limitations. First, the approaches used here do not provide mechanistic insights into the process of phase separation, that is, whether condensate formation is driven purely by liquid-liquid <u>p</u>hase <u>s</u>eparation (LLPS), phase separation coupled to percolation (PSCP), or phase separation coupled to other phase transitions (PS++)^9^. The current framework also does not include an assessment of the boundary between the dense and dilute phases, which is important in understanding the finite interfacial tension that hinders macromolecular transport across the boundary. Also, for broad use and accessibility, we developed the experimental framework using heterologous protein expression in *E. coli*, which may not accurately reflect the *in vivo* conditions of the native host. Finally, low-level leaky expression of inducible promoters may preclude the examination of proteins that phase separate at very low *in vivo c*_sat_. Despite these limitations, this framework provides broad-use and systematic approaches that address ongoing debates over the rigor and standardization of phase separation assessments in bacterial cells.

## Methods

### Bacterial strains, plasmids, and primers

Strains, plasmids, and primers used in this study are listed in Tables S1 and S2. All constructs were made using Gibson assembly^40^ from PCR fragments or synthesized dsDNA (Integrated DNA Technologies) and verified by Sanger sequencing. For example, plasmid pCA3 was generated from plasmid pTrc99A-mCherry-*cI78^EP^* by replacing mCherry with PAmCherry. The PAmCherry fragment was generated using primers YH1 F and YH1 R. The plasmid pTrc99A-mCherry-*cI78^EP8^* was amplified using YH2 F and YH2 R to generate the second fragment. The two fragments were then added to a Gibson assembly reaction to enzymatically join the overlapping DNA sequences. Other plasmids were generated using similar methods with primers indicated in Table S2. When relevant, homology regions for Gibson assembly are indicated in blue. Plasmids were introduced into their respective host strains by chemical transformation and selection for antibiotic resistance encoded by the plasmid. All plasmids are available on AddGene.

### Growth conditions

Lysogeny broth (LB) and AB media were used as either a broth or solid for culturing bacteria. LB medium was used to grow *E. coli* BL21 <u>A</u>rctic <u>E</u>xpress (AE) and overnight cultures of *E. coli* MG1665. The minimal AB medium was used when inducing protein expression in *E. coli* MG1665 to ensure protein expression reproducibility, as all the components are defined; as opposed to a complex medium such as LB^28,41^. AB medium was supplemented with 0.2% of a carbon source (glycerol for growth or glucose to also inhibit basal protein expression from the P*_trc_* promoter), 0.2% casamino acids, 10 μg/ml thiamine, and 25 μg/ml uracil^28^. *E*. *coli* was grown in a 15-ml tube overnight for approximately 15 h in 5 ml of LB broth at 37 °C on an orbital shaker at 200–225 rpm. Exponential phase cultures were prepared by diluting overnight cultures 1:100 and further incubated until an OD_600_ of 0.2–0.6 was reached. When appropriate, the following chemicals were added to the medium at the indicated final concentrations: carbenicillin (100 μg/ml) for selecting the plasmids in culture, IPTG (0.1–1 mM) or L-arabinose (0.2%) for protein induction.

### Total protein and immunoblot analyses

A 0.4-ml aliquot of *E. coli* cells (OD_600_: 0.2–0.4) was lysed using a Qsonica cupped-horn sonication system (20 cycles, 30 s on, 10 s off at 30% power) and centrifuged at 10,000 x g for 1 min at 4 °C. The protein content in the supernatant was measured using a Bradford assay kit according to the manufacturer’s instructions. To prepare samples for immunoblot analysis, an equal volume of 4x Laemmli sample buffer was added to *E. coli* culture prior to boiling 20 min. One microgram of total protein from each sample was loaded in each lane of a 4-12% Bis-Tris NuPAGE gel. Gels were transferred onto a mini-size polyvinylidene difluoride membrane using a Trans-Blot Turbo system (Bio-Rad). The membrane was immunoprobed using a rabbit polyclonal antiserum against mCherry (1:2000). The membrane was then incubated with the goat anti-rabbit IgG Secondary Antibody IRDye 800CW. Membrane signals were visualized and quantified using LI-COR Image Studio. The mCherry band of each lane was normalized to the total intensity of the lane to calculate the degradation level of mCherry fusion proteins.

### Wide-field fluorescence and phase-contrast imaging

Wide-field fluorescence and phase-contrast imaging were performed using a Nikon Ti2-E motorized inverted microscope controlled by the NIS Elements software with a SOLA 365 LED light source, a 100× objective lens (Oil CFI Plan Apochromat DM Lambda Series for Phase Contrast), and a Photometrics Prime 95B back-illuminated sCMOS camera or Hamamatsu Orca-Flash 4.0 LTS camera. Fusions to mNeonGreen (mNG) were imaged using a “YFP” filter set (C-FL YFP, Hard Coat, High Signal-to-Noise, Zero Shift, Excitation: 500±10 nm, Emission: 535±15 nm, Dichroic Mirror: 515 nm). Fusions to msfGFP were imaged using a “GFP” filter set (C-FL GFP, Hard Coat, High Signal-to-Noise, Zero Shift, Excitation: 436±10 nm, Emission: 480±20 nm, Dichroic Mirror: 455 nm). DAPI fluorescence was imaged using a “DAPI” filter set (C-FL DAPI, Hard Coat, High Signal-to-Noise, Zero Shift, Excitation: 350±25 nm, Emission: 460±25 nm, Dichroic Mirror: 400 nm). mCherry was imaged using a “Texas Red” filter set (C-FL Texas Red, Hard Coat, High Signal-to-Noise, Zero Shift, Excitation: 560±20 nm, Emission: 630±37.5 nm; Dichroic Mirror: 585 nm).

### Time-lapse videos of protein induction in *E. coli BL21*

Gene *mNG-McdB* and *mNG-cI^agg^* were cloned into the multiple cloning site of the vector pET11a to create the pJB37 and pYH73 plasmids respectively, used for inducible expression under the control of a bacteriophage T7 promoter. pJB37and pYH73 were transformed into BL21 (AE) cells and a 5 ml overnight culture containing 100 μg/ml of carbenicillin in LB medium was grown at 37 °C with shaking at 225 rpm. The overnight culture (50 μl) was used to inoculate 5 ml of LB supplemented with 100 μg/ml of carbenicillin in a 15-ml tube. The cells were grown at 37 °C with shaking at 225 rpm to an OD_600_ of 0.2–0.6. Protein expression was then induced by the addition of 1 mM IPTG and 0.2% Arabinose solution to the tube. Cells (2 µl) were spotted on a 1 cm diameter pad made of 1.5% UltraPure agarose in LB and supplemented with 1mM IPTG/0.2% Arabinose. After two minutes, the cell-containing side of the pad was flipped onto a 35-mm glass-bottom dish and mounted onto the stage of a Nikon Ti2-E motorized inverted microscope controlled by NIS Elements software with a SOLA 365 LED light source, a 100x objective lens (Oil CFI Plan Apochromat DM Lambda Series for Phase Contrast), and a Photometrics Prime 95B back-illuminated sCMOS camera. mNG was imaged using the “YFP” filter set (see above). An image series was captured every 1 min for 4 hours.

### DAPI staining

*E. coli* BL21 (AE) cells with plasmid pJB37 were induced with 1mM IPTG/0.2% Arabinose as described in the “Time-lapse videos of protein induction in *E. coli* BL21” section. DAPI was added to the exponentially growing culture at a final concentration of 2 μM. Cells were incubated in DAPI for 15 min at 25° C before imaging, without rinsing. DAPI was imaged using the “DAPI” filter set (see above).

### Localized cell lysis

pJB37 was transformed into BL21 (AE) cells, and a 10 ml overnight culture containing 100 μg/ml of carbenicillin was grown at 20 °C with shaking at 225 rpm. The overnight culture (0.5 ml) was used to inoculate 50 ml of LB supplemented with 100 μg/ml of carbenicillin in a 250 ml baffled flask. The cells were grown at 37 °C with shaking at 225 rpm to an optical density of 0.5. The flasks were plunged into an ice bath for two minutes. Protein expression was then induced by the addition of 1 mM IPTG/0.2% Arabinose solution to the flask. Flasks were then returned to the shaker with incubation temperature set to 16 °C. Cells were then grown overnight with shaking at 225 rpm (∼16 h induction). Cells (2 µl) were spotted on a 1 cm diameter pad made of 1.5% UltraPure agarose in LB. After two minutes, the cell-containing side of the pad was flipped onto a 35-mm glass-bottom dish and mounted onto the stage of a Nikon Ti2-E motorized inverted microscope controlled by NIS Elements software with a SOLA 365 LED light source, a 100x objective lens (Oil CFI Plan Apochromat DM Lambda Series for Phase Contrast), and a Photometrics Prime 95B back-illuminated sCMOS camera. mNG-McdB was imaged using the “YFP” filter set (see above). A region of interest (0.5 µm diameter) was drawn at one cell pole and used as the target for a 405 nm laser pulse (1 sec at 50 mW), which caused localized cell lysis. An image series was captured every 500 ms for 60 seconds.

### Temperature shift

*E. coli* BL21 (AE) cells with plasmid pJB37 were grown to an OD of 0.5 and then flasks were plunged in an ice bath as described above for 2 min. mNG-McdB expression was then induced by the addition of 1 mM IPTG/0.2% Arabinose solution to the flask. Cells (2 µl) were then immediately spotted on a 1 cm diameter agarose pad. After two minutes the cell-containing side of the pad was flipped onto a 35-mm glass-bottom dish and mounted onto a stage-top incubator with temperature control. mNG-McdB expression and focus formation was imaged using the “YFP” filter set (see above). Once mNG-McdB initiated focus formation, the stage-top temperature was ramped up from 25 to 37 °C. An image series was captured every 5 s for 30 min to observe focus dissolution.

### Cellular shape change in *E. coli* BL21

*E. coli* BL21 (AE) cells with plasmid pJB37 and pYH73 were induced with 1mM IPTG/0.2% Arabinose as described in the “Time-lapse videos of protein induction in *E. coli* BL21” section. The MreB inhibitor A22 (10 µg/ml) was added to the exponential phase cultures at time *t*=0. The cultures were then incubated at 37 °C with shaking at 225 rpm for 6 hours. Images were taken at 6 h post-treatment with the “YFP” filter set (see the “Wide-field fluorescence and phase-contrast imaging” section).

### Tunable induction of mCherry fusion focus

Gene *mCherry-cI^agg^, mCherry-PopTag^SL^, mCherry-PopTag^LL^, mCherry-mcdB, mCherry-mcdB*^sol^, and *mCherry* were cloned into the multiple cloning site of the vector pTrc99A to create the pTrc99A-*mCherry-cI78^EP8^,* pYH75, pYH80, pYH71, pYH86, and pYH77 plasmids respectively. The above plasmids were transformed into MG1665 cells and a 5 ml overnight culture containing 100 μg/ml of carbenicillin in AB medium (with 0.2% glycerol as the carbon source) was grown at 37 °C with shaking at 200 rpm. The overnight culture (50 μl) was used to inoculate 5 ml of AB (with 0.2% glycerol as the carbon source) supplemented with 100 μg/ml of carbenicillin in a 15-ml tube. The cells were grown at 37 °C with shaking at 200 rpm to an OD_600_ of 0.2–0.6. 500 μM of IPTG was added to the exponential phase culture to induce fluorescent fusion protein expression. After each hour, 2 μL of the MG1665 culture were spotted on an agarose round pad. The pads were prepared by dissolving Ultrapure agarose in AB to a final concentration of 1.5%. A series of images were taken with the “Texas Red” filter set (see the “Wide-field fluorescence and phase-contrast imaging” section) every 1 hour for 5 hours.

### Nucleoid compaction

*E. coli MG1665* with pYH86 plasmid was induced with 1 mM IPTG for 2 hours. Cells were then stained with 2 μM DAPI and spotted on an agarose pad that has 50 μM of ciprofloxacin. A series of images were taken with the “Texas Red” and “DAPI” filter sets (see the “Wide-field fluorescence and phase-contrast imaging” section) every 15 min for 7 hours.

### Time-lapse videos that examined focus reversibility in *E. coli* MG1665

*E. coli MG1665* with pTrc99A-*mCherry-cI78^EP8^,* pYH75, pYH80, pYH71, pYH86, and pYH77 plasmids were induced with 1mM IPTG for 2 h. The cells were then washed three times with 10X volume of fresh AB media supplemented with 0.2% glucose and incubated for 30 min at room temperature before imaging. After 30 min, 2 μL of the MG1665 culture were spotted on an agarose round pad. The pads were prepared by dissolving Ultrapure agarose in AB to a final concentration of 1.5% (the AB medium contains 0.2% glucose and 10 μg/ml cephalexin when specified). A series of images were taken with the “Texas Red” filter set (see the “Wide-field fluorescence and phase-contrast imaging” section) every 15 min for 15 hours.

### Fluorescence recovery after photobleaching (FRAP)

*E. coli MG1665* with pTrc99A-*mCherry-cI78^EP8^,* pYH75, pYH80, and pYH71 plasmids were induced with 1mM IPTG for 2 h. FRAP was performed after cells were washed with fresh AB media supplemented with 0.2% glucose and incubated for 30 min. A Nikon Ti2-E motorized inverted microscope controlled by NIS Elements software with a SOLA 365 LED light source, a 100x objective lens (Oil CFI Plan Apochromat DM Lambda Series for Phase Contrast), and a Photometrics Prime 95B back-illuminated sCMOS camera were used to do FRAP experiments. Control images were taken before bleaching, then the regions of interest (foci) were bleached with a laser at 405 nm and 30% power (15 mW). A series of images was captured every 500 ms for 60 seconds after bleaching. More than ten different regions of interest (ROIs) were chosen per sample. Image analysis was performed using ImageJ. The background signal was subtracted from the bleached focus and an unbleached focus within the cell. The resulting focus signal was normalized such that the pre-bleach signal is one and the first frame post-bleaching is zero.

### Imaging the IbpA protein with wide-field fluorescence microscopy

The plasmids pTrc99A-*mCherry-cI78^EP8^,* pYH75, pYH80, pYH71, and pYH77 plasmids were transformed into *E. coli* MG1665 that expresses IbpA-msfGFP by its native promoter. A 5 ml overnight culture containing 100 μg/ml of carbenicillin in AB medium (with 0.2% glycerol as the carbon source) of each strain was grown at 37 °C with shaking at 200 rpm. The overnight culture (50 μl) was used to inoculate 5 ml of AB (with 0.2% glycerol as the carbon source) supplemented with 100 μg/ml of carbenicillin in a 15-ml tube. The cells were grown at 37 °C with shaking at 200 rpm to an OD_600_ of 0.2–0.6. 1mM of IPTG was added to the exponential phase culture to induce fluorescent fusion protein expression for 2 h. 2 μL of the MG1665 cultures were spotted on an agarose round pad. The pads were prepared by dissolving Ultrapure agarose in AB to a final concentration of 1.5%. Images were taken with the “GFP” and “Texas Red” filter set (see the “Wide-field fluorescence and phase-contrast imaging” section) at 2 h post-induction.

### Imaging the IbpA protein with 2D Structured illumination microscopy (SIM)

The agarose pads with induced MG1665 cells were prepared similar to those in the “Imaging the IbpA protein with wide-field fluorescence microscopy” section. SIM images were acquired on a Nikon N-SIM system equipped with a Nikon SR HP Apo TIRF 100X 1.49NA objective, a Hamamatsu ORCA-Flash4.0 camera (65 nm per pixel), and 488 nm and 561 nm lasers from a Nikon LU-NV laser launch. Cells were identified using DIC to avoid photobleaching. For each 2D-SIM image, nine images were acquired in different phases via the built-in 2D SIM modes. Super-resolution image reconstruction was performed using the Nikon Elements SIM module.

### Single-molecule fluorescence microscopy

Cells expressing protein fusions to PAmCherry of a plasmid with the pTrc inducible promoter were grown in 3 ml of lysogeny broth (LB) medium in a culture tube for ∼ 14 h, with shaking at 225 rpm at 37 °C. The following day, cells were diluted 1:100 into 3 ml of fresh AB medium supplemented with 0.2% glucose or glycerol and grown to OD_600_ ∼ 0.3 before imaging either as is (“no induction” condition) or inducing with 100 µM IPTG (“inducing” condition). After a two- or four-hour induction, cells were washed three times with 1 ml of AB medium supplemented with 0.2% glucose. All “no induction” and “inducing” cells were resuspended in M9 medium supplemented with 0.2% glucose for imaging. Agarose pads were made at 2% (w/v) with M9 minimal medium supplemented with 0.2% glucose. An aliquot of 2.5 µL of cells was loaded onto an agarose pad and sandwiched between two coverslips. Cells were imaged at room temperature with a 100× 1.40 numerical aperture oil-immersion objective. A 406-nm laser (Coherent Cube 405-100; 0.2 W/cm^2^) was used for PAmCherry photoactivation and a 561-nm laser (Coherent-Sapphire 561-50: 88.4 W/cm^2^) was used for imaging. Given the difference in protein expression level, 200 – 400 ms activation doses were used for cells not induced and 50 ms activation doses were used for the two hour induction sample. The four hour sample had many preactivated molecules and molecules activated spontaneously without activation via 405-nm laser, which limited our ability to confidently localize single molecules. To reduce the number of molecules per imaging frame, an initial 405-nm laser activation (10 – 15 s) was followed by photobleaching through illumination with a 561-nm laser with the same power density as above for 15-20 minutes or until spatially resolved single molecules were observed. The fluorescence emission was filtered to eliminate the 561-nm excitation source and imaged at a rate of 40 ms/frame using a 512 x 512-pixel Photometric Evolve EMCCD camera.

### Single-molecule data analysis

Phase-contrast microscopy images were used to segment bacterial cells (see *Cell segmentation* for details) prior to localization and tracking. Single molecules were detected and localized with a 2D Gaussian fitting by the SMALL-LABS algorithm^42^ and connected into trajectories using the Hungarian algorithm^43^. Localization heat maps (Fig. 2f) were made by normalizing the segmented cells and rotating them onto their long axes, followed by projection, binning, and symmetrization of the single-molecule localizations onto the normalized cell^44,45^.

Only single-molecule trajectories with a minimum of six steps were used for trajectory analysis. To extract the apparent diffusion coefficient for each trajectory, a modified version of the diffusion model^46^ was fit to the mean square displacement (MSD) as a function of time lag over the time interval 40 ≤ *τ* ≤ 200 ms. The modified diffusion model was used to account for motion blur due to averaging the true position of a molecule over the time of a single acquisition frame^46^.

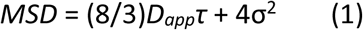

In this case, *D_app_* is the apparent diffusion coefficient, *τ* is the time lag, and σ describes the localization precision. Only fits with *R*^2^ ≥ 0.7 were kept. The log distribution histograms of the *D_app_* were each fit to a two-state Gaussian mixture to determine the *D_app_* and the associated weight fraction for the slow and fast diffusion modes (Supplementary Fig. 7).

### Measuring single-cell protein concentrations

To determine the average number of photons detected from a single mCherry molecule per imaging frame, cells expressing the indicated mCherry fusion proteins were grown in 3 ml of AB medium supplemented with 0.2% glucose to OD ∼0.3 and washed once in 1 ml of M9 minimal medium prior to imaging. Cells were pre-bleached with a 561-nm laser until only a few isolated molecules were observed. Images were then recorded at 40-ms exposure with a 561-nm laser (110.5 W/cm^2^) and an input EM gain of 600 (NIS-Elements software setting). Single molecules were detected as described earlier. The number of photons detected per single molecule per imaging frame was obtained by calculating the integrated intensity counts of the 2D Gaussian fit from the localization step^47^, which was then converted into the number of photons using the following camera calibrations.

The conversion gain (number of photoelectrons per fluorescence intensity count) calibration was performed as described in a Teledyne Photometrics Technical Note^48^. Briefly, multiple images of a white business card were acquired for 10-, 20-, 40-, 80-, 160-, and 320-ms exposure times with no electron multiplication (EM) gain. To account for the camera bias, 100 frames were acquired with nominally 0-ms exposure and a shuttered camera path. The average of these 100 frames was subtracted from every subsequently analyzed image. A plot of the mean signal of the image for each exposure time versus the variance of the same image gives a straight line with a slope that equals the conversion gain (Supplementary Fig. 3a). Pixel-to-pixel nonuniformity effects were removed by recording two images at each exposure time^49^. The conversion gain of our camera was found to be 1.40 electrons per intensity count.

To determine the EM gain (the number of electrons per photoelectron), two images were recorded: one long-exposure image (1 s) with no EM gain and one short-exposure image (10 ms) with an arbitrary EM gain. The bias was subtracted from both images. The EM gain multiplication factor is the factor difference in signal per time unit between the corrected images. We repeated this procedure for various software setting EM gains and found a linear fit in the range of 5–600 input EM gain with a conversion factor of 0.15 between nominal EM gain and output EM gain (Supplementary Fig. 3b).

To calculate the number of photons from the fluorescence image, we therefore multiplied the intensity counts recorded by the conversion gain of 1.40 and divided this quantity by the input EM gain multiplied by the EM gain conversion factor of 0.15. The distribution of photons per molecule per imaging frame was fit to a Gamma distribution resulting in a peak of ∼90 photons per molecule per imaging frame (Supplementary Fig. 3c).

To determine the apparent cellular concentration of mCherry-McdB and mCherry-PopTag^LL^ when foci are present, cells expressing these proteins were grown and prepared as described above. The mCherry-McdB strain was induced with 1 mM IPTG for 2 h prior to imaging and the mCherry-PopTag^LL^ was induced with 100 µM IPTG for 2 h prior to imaging. Cells were imaged with a 561-nm laser (110.5 W/cm^2^) at 20 ms exposure and 10x (software setting) EM gain. To minimize the effect of photobleaching on the photon counting measurement, image acquisition was started prior to laser illumination.

The brightest five frames in the movie were averaged to determine the cell brightness before photobleaching. The integrated total fluorescence emission within a cell is linearly related to the total quantity of McdB or PopTag^LL^ molecules per cell: this intensity value was divided by the number of photons per mCherry molecule per imaging frame to obtain the McdB or PopTag^LL^ copy number per cell. To determine the cellular concentration of the proteins, the volume of the cell was estimated by modeling it as a cylinder with spherical caps^50^.

### Image analysis

#### Cell segmentation

Cell segmentation was performed with the Cellpose^51^ package in Python. To train the model optimally for bacterial cell morphology, 26 raw phase contrast images of cells were manually annotated. To segment the cells using the trained model, a Gaussian blur (standard deviation of Gaussian = 0.066 µm) was applied to the bacterial cell phase-contrast images and the blurred cells were segmented. Cells touching the borders of the image were ignored. Erroneous segmentations were manually corrected using the Cellpose GUI or excluded from further analysis.

#### Condensation analysis

Condensation coefficients were calculated as described previously^16,31,52^. Briefly, the fluorescence intensity for each pixel in a cell was corrected for background by subtracting the median value of all pixels in an image outside of the cell regions and normalized by the minimum and maximum pixel intensity values in the corresponding cell:

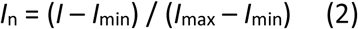

These normalized pixel intensities were then binned to generate histograms that represent the localization pattern for each protein (Supplementary Fig. 4a-f).

Next, cells were classified by the presence or lack of a focus. An ROI was defined for each cell using the segmentation phase mask. Each ROI was applied a Gaussian blur (SD = 0.066 µm) and normalized using Eq. (2). Next, foci were detected by generating a binary image where only pixels above a specified intensity threshold were assigned a non-zero value. The putative foci were then filtered by area and eccentricity (see Supplementary Table 5 for parameters used). Homogeneously distributed protein mCherry or cells without a detected focus display a flat distribution whereas the strongly clustered protein cI^agg^ displays a strongly left-skewed distribution (Supplementary Fig. 4a-f). To quantify the differences between the proteins measured, we calculated the fraction of pixels with a normalized intensity below a threshold value, *I* < 0.3, 0.5, and 0.7, for each cell (Supplementary Fig. 4g). These values were then normalized to the condensation coefficient distribution of mCherry as our experimental representative of a homogenous distribution.

#### Partition ratio analysis

Foci partition ratios were calculated from cells with a detected focus (see *Condensation analysis*) as the ratio of the average background-corrected pixel intensity within a focus to the average background-corrected pixel intensity of the cells excluding the focus region. For cells with two or more detected foci, the average intensity of all foci was used for the ratio.

#### Focus dissolution analysis

Fluorescent foci were tracked during the reversibility experiment for images acquired over the course of bacterial cell division events or during cell elongation with cephalexin treatment as described above. We detected fluorescent foci by the LoG method in TrackMate^53^ using an estimated blob diameter of 600 nm and a quality threshold of 80 for cI^agg^, PopTag^SL^, PopTag^LL^, and 225 for McdB. Spot detections were subsequently linked to generate trajectories using the Simple Lap Track option with a maximum linking distance of three pixels (198 nm), a maximum gap-closing distance of 3 pixels (198 nm), and a maximum gap of 2 frames (30 min). The following trajectory filters were used: (i) only trajectories present in the first frame were tracked in subsequent frames, (ii) trajectories of foci in cells that move out of the field of view during the movie were excluded, (iii) false positive detections in the first frame were removed, and (iv) trajectories of foci in cells that did not grow (cephalexin treatment) or divide (generational dilution) were excluded.

#### IbpA association analysis

Images of dual-color labeled cells expressing msGFP-IbpA and mCherry fusion proteins were analyzed by detecting fluorescent foci in the mCherry channel using an intensity-based threshold of 30% of the maximum intensity in the image. mCherry fusion protein spot detections were subsequently filtered based on a minimum area threshold of 4 pixels and an eccentricity threshold of 0.75. The centroid of these spots was used to define the center of a 23 x 23 pixel ROI. This ROI was then used to analyze the IbpA channel; first an intensity profile was collected using the centroid of the ROI as the center point for both channels and subsequently normalized to the max intensity values in the mCherry channel. Next, the normalized ROIs for each channel were averaged to generate the projection images (See Figure 5c). The resulting intensity patterns for both channels were then fit with a 2D Gaussian to measure the full width at half max (FWHM) of the intensity. Finally, we calculated the ratio of FWHM_IbpA_/FWHM_mCherry_.

## Data Availability

All data generated or analyzed during this study are included in this published article [and its supplementary information files]. Source data are also provided with this paper.

## Code Availability

Data processing and analysis scripts for this study were written in MATLAB and Python. All image analysis scripts, and the SMALL-LABS and NOBIAS algorithm packages are available on GitHub (https://github.com/BiteenMatlab).

## Acknowledgements

This work was funded by NSF CAREER award 1941966 to AGV and NIH award R01GM144731 to JSB. We would like to thank Dr. Sarah Veatch for providing critical feedback in experimental direction and data interpretation. We also thank Dr. Eric Rentchler of the Biomedical Microscopy Core at the University of Michigan for training and coordinating access to 2D-SIM. We thank Dr. Ming Li’s Lab at the University of Michigan for the mCherry antibody, Dr. Abram Aertsen’s Lab for sending plasmids and *E. coli* strains. We thank Dr. Julien Mortier for constructing the strain that expressed IbpA-msfGFP with its native promoter in *E. coli*.

---

**Figure S1.**
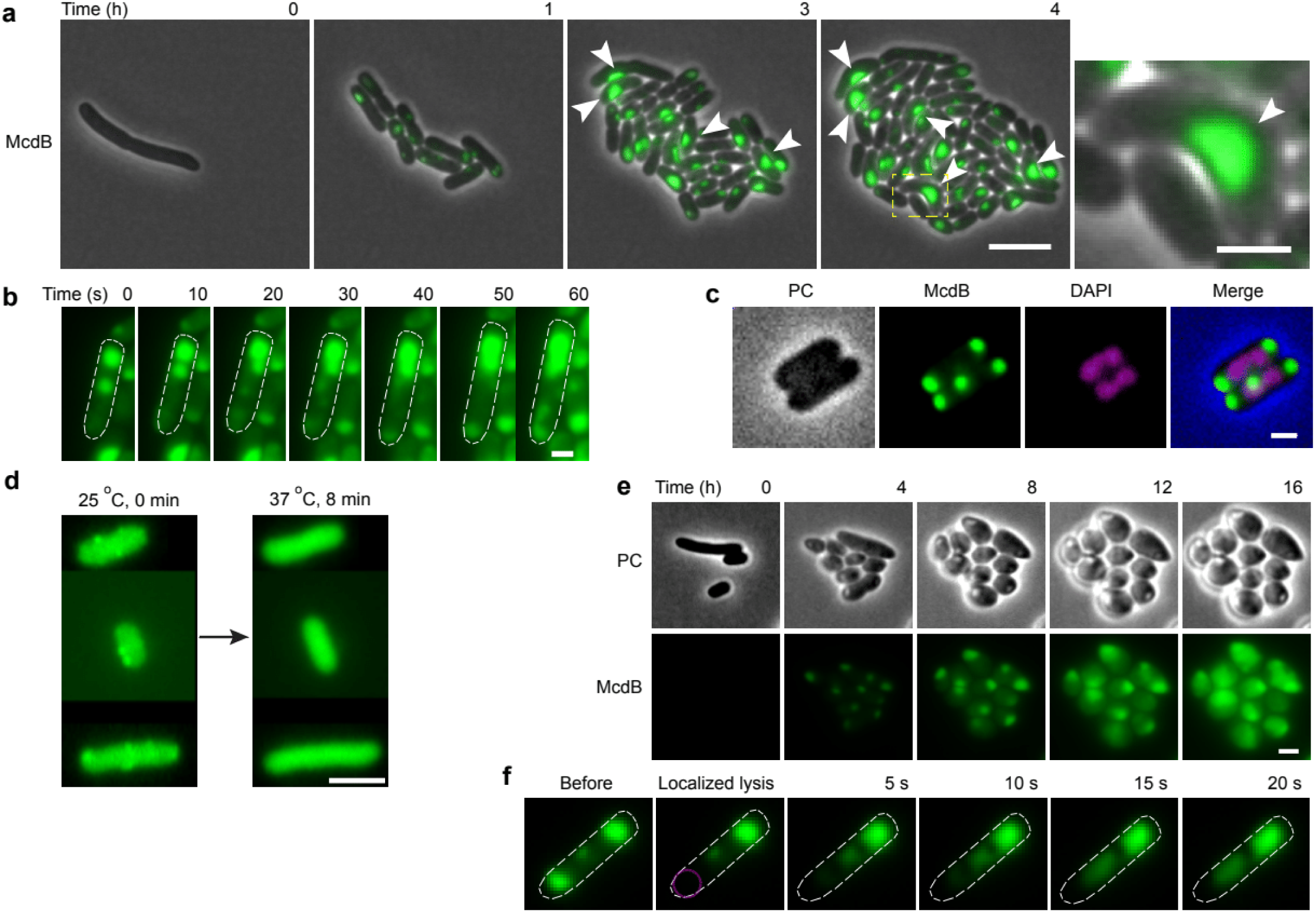
Overexpression of mNG-McdB in *E. coli* BL21. **a**. mNG-McdB forms foci. The phase contrast channel is shown in gray and the mNG channel is green. The right-most image is a zoom-in of the area with the dashed yellow box. White arrows highlight cells where McdB foci wet to the inner membrane inducing cell curvature. Images are representative of three biological replicates. Left scale bar: 5 µm; Right scale bar: 2 µm. **b**. Fusion of mNG-McdB foci. Representative frames show two foci merging together in 60 s. Scale bar: 1 µm. **c**. mNG-McdB foci are nucleoid-excluded. DAPI stain is shown in magenta. Scale bar: 1 µm. **d**. Effects of temperature shift on focus stability. Cells that formed foci at 25 °C were transitioned to 37 °C in a stage-top incubator. Representative frames at 0 and 8 min are shown. Scale bar: 1 µm. **e**. Effects of changing cell volume on focus stability. *E. coli* cell volume was increased by treating cells with the MreB inhibitor, A22 (10 µg/ml). Scale bar: 2 µm. **f**. Localized lysis of the cell. One cell pole was lysed using a high intensity laser, focused within the indicated magenta circle. Representative frames show solubilization of the opposing focus and shift of fluorescent signal to the opposite end of the cell. Scale bar: 1 µm.

**Figure S2.**
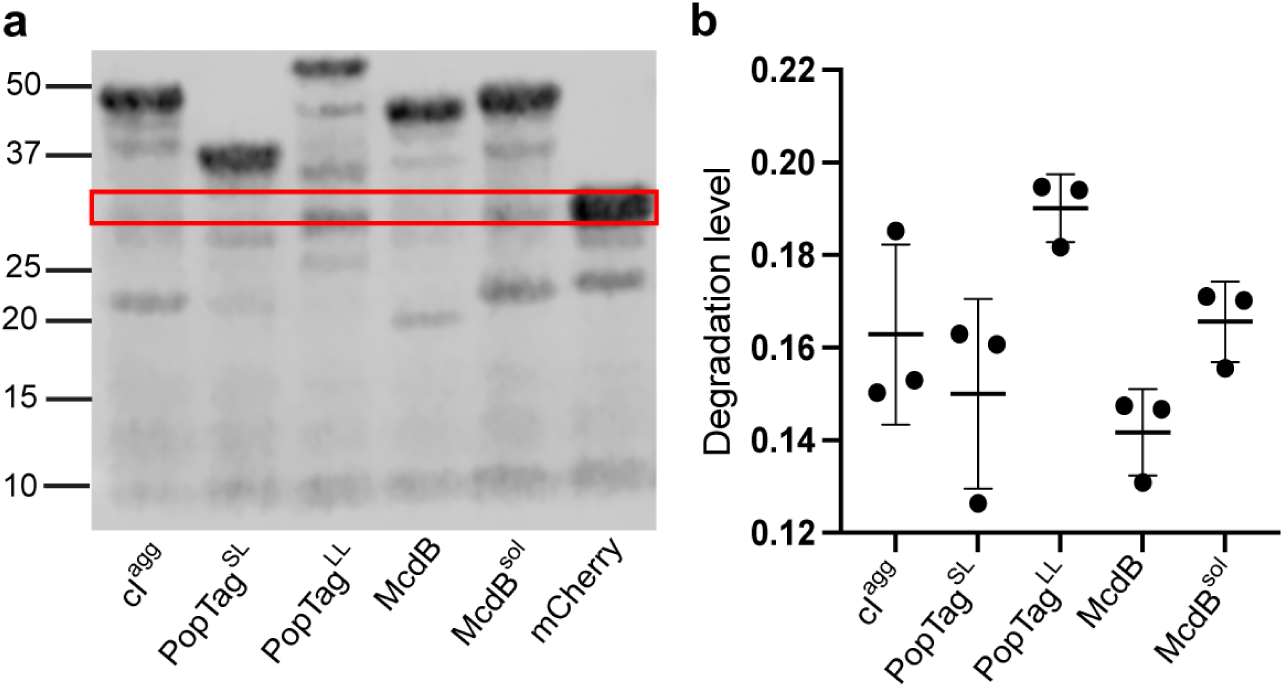
Expression of mCherry fusion proteins. *E. coli* MG1665 cells of indicated strains were induced with 1 mM IPTG for 2 h. **a**. Immunoblot analysis of mCherry fusion proteins. Representative immunoblots are shown. Red box shows the band at the mCherry size (28 kDa). Similar results were observed in three biological replicates. **b**. Degradation level of mCherry fusion proteins. In each lane, the intensity of the band at the mCherry size was quantified and normalized to the total intensity of the respective lane. Error bars represent the standard deviation from three biological replicates.

**Figure S3.**
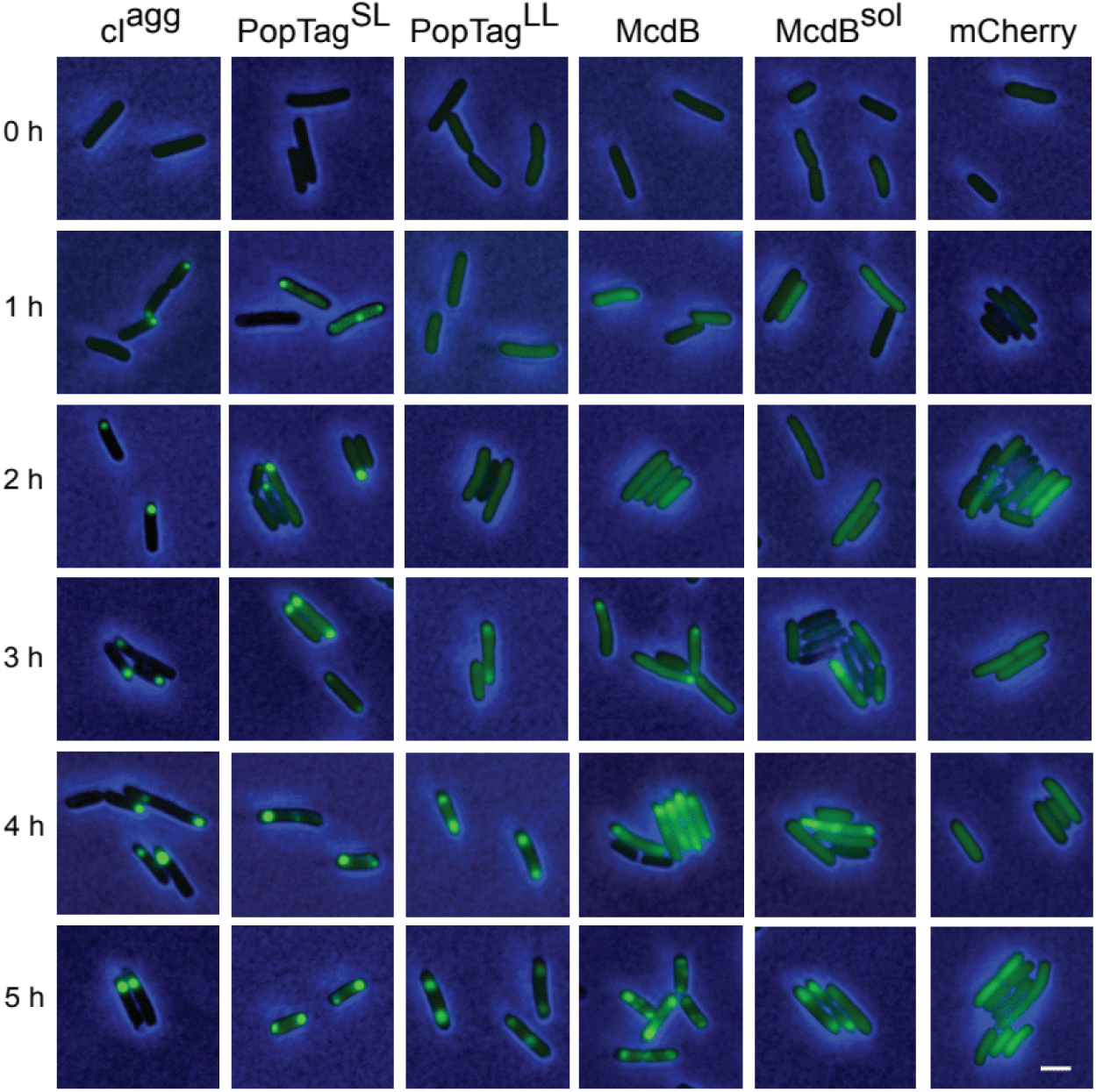
Expression of mCherry fusion proteins. Representative images of indicated mCherry fusion protein foci in *E. coli* from 0 to 5 h post-induction. Phase contrast (blue) and mCherry (green) channels were merged. Images are representative of three biological replicates. Scale bar: 1 µm.

**Figure S4.**
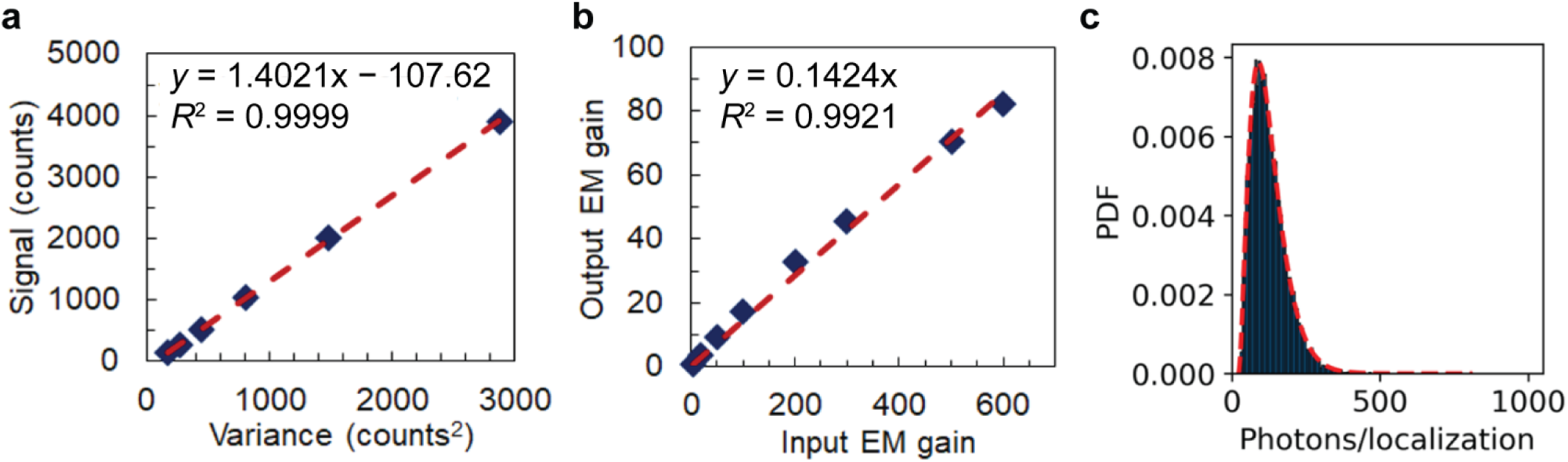
Electron-multiplying charge coupled detector (EMCCD) calibration for photon counting. **a**. Signal variance versus signal average for different camera integration times. The data points correspond to the following camera integration times: 10, 20, 40, 80, 160, and 320 ms. The dashed red line is the linear regression fit. **b**. Electron multiplication conversion plot. The dashed red line is the linear regression fit. **c**. Distribution of number of detected photons per localization per imaging frame for mCherry-McdB. Signal histogram was fit to a gamma distribution resulting in a peak of 91.1 photons per localization at a 40-ms camera integration time. 35800 localizations were used to build the histogram.

**Figure S5.**
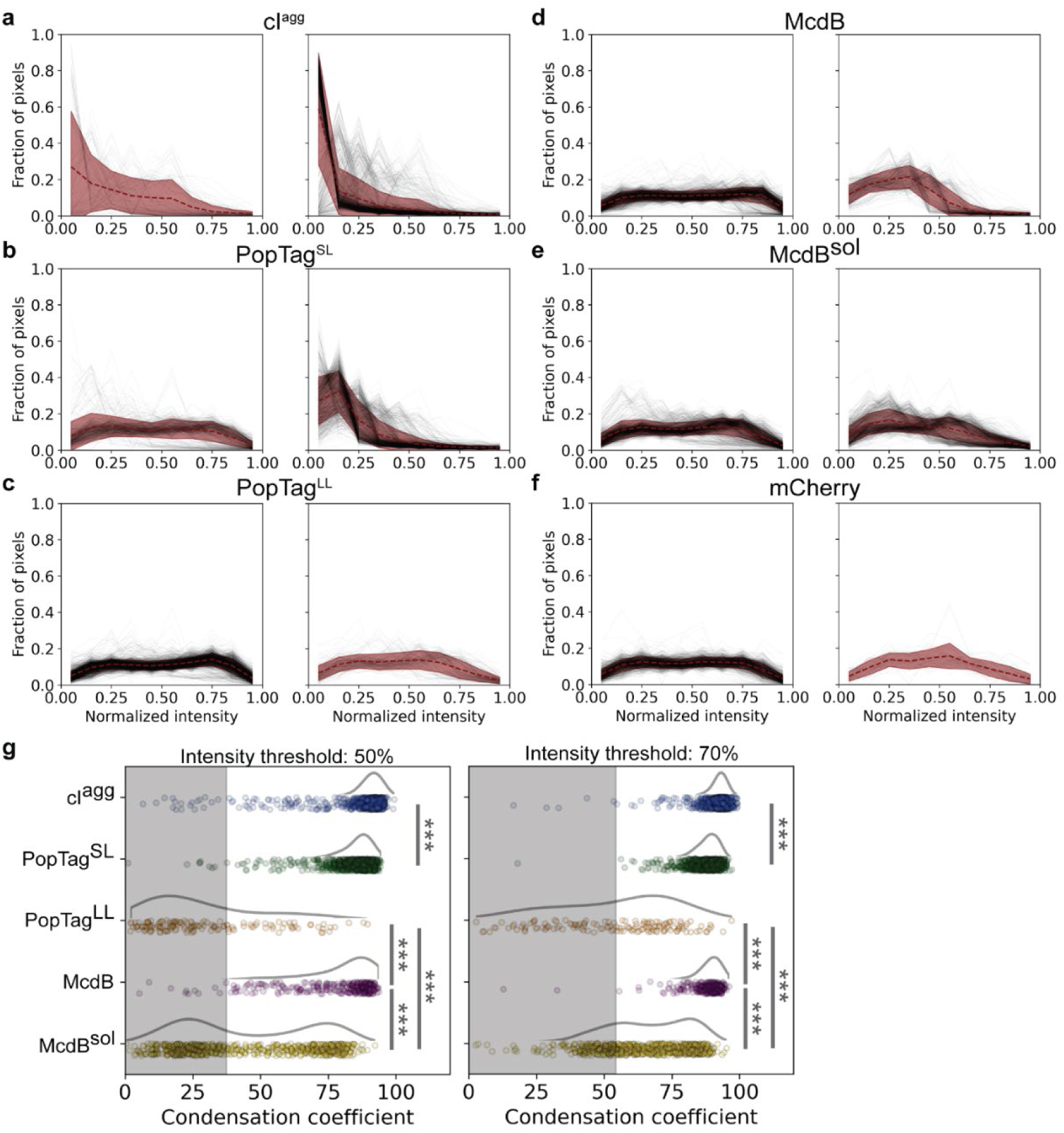
Normalized pixel intensity histograms and condensation coefficients of mCherry fusion proteins. **a-f**. Normalized pixel intensity histograms for cells with no detected focus (left) and a detected focus (right). Black lines represent individual cells. The red dashed line and shading are the average and standard deviation across cells for each plot. **g**. Quantification of condensation coefficients. Condensation coefficient plots for cells with a detected focus and threshold values of 50 and 70 % of the max intensity. Data points correspond to individual cells. The curves next to the scatter plots were obtained via kernel density estimation. The shaded region represents the measurement range for cells expressing a uniform mCherry signal. Analysis was done on *N* > 900 cells for each fusion over three biological replicates.

**Figure S6.**
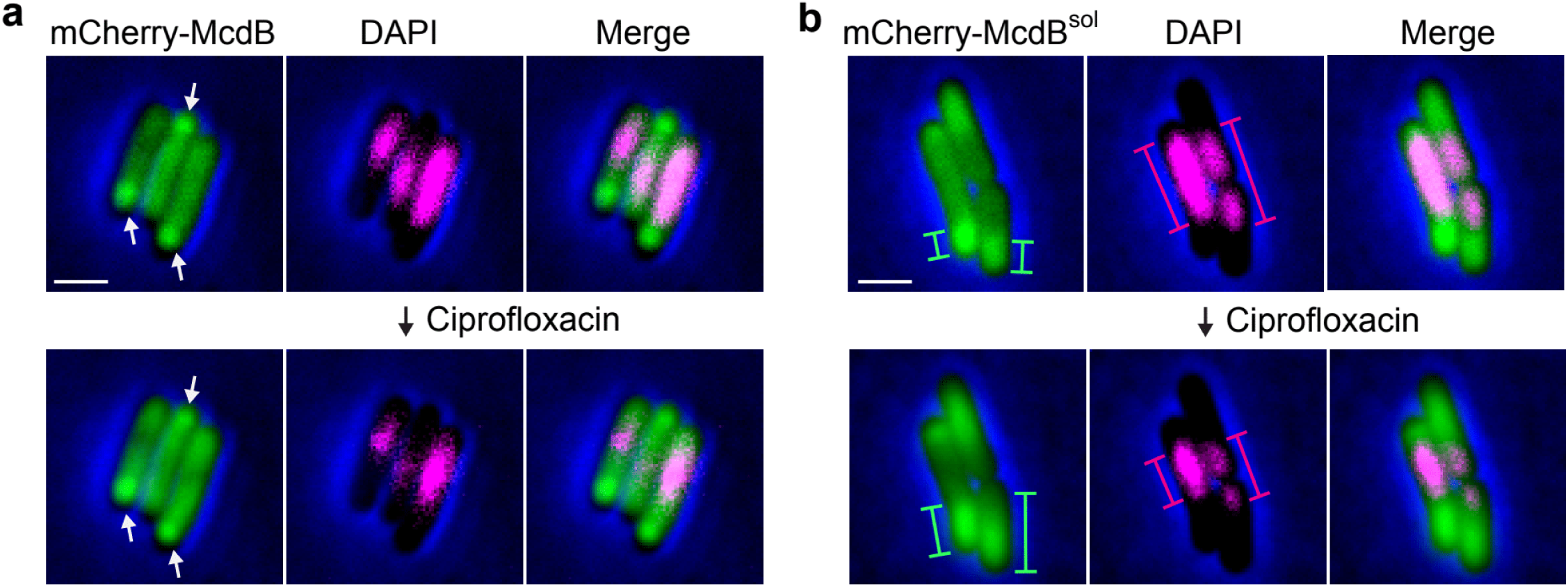
mCherry-McdB^sol^ condenses by repulsive interactions with the nucleoid. Representative images of mCherry-McdB (**a**) and mCherry-McdB^sol^ (**b**) foci in *E. coli* at 4 h post-induction (top). Cells were then treated with ciprofloxacin (50 mM) for nucleoid compaction and stained with DAPI (2 µM) for nucleoid visualization (bottom). mCherry (green), DAPI (magenta), and merged images overlaid with Phase contrast (blue) are shown. White arrows highlight the mCherry-McdB foci, which remained after nucleoid compaction. Green and magenta bars indicate the length of the mCherry-McdB^sol^ foci and the nucleoid respectively before and after ciprofloxacin treatment. As the nucleoid condensed, mCherry-McdB^sol^ foci expanded. Images are representative of three biological replicates. Scale bars: 1 µm.

**Figure S7.**
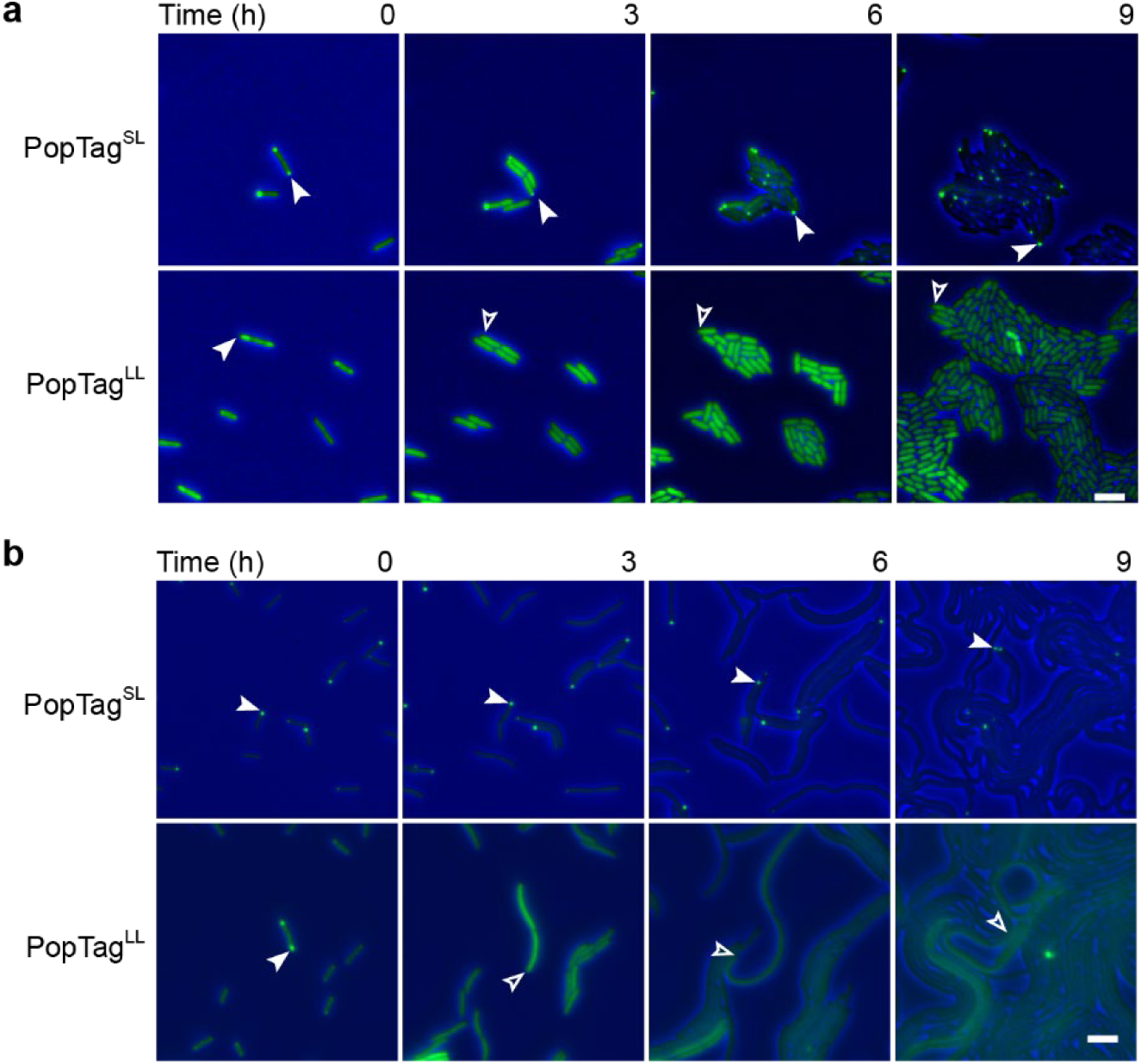
Cell growth and division dissolve foci by dropping the cellular concentration below *c*_sat_app_. **a.** Generational-dilution dissolves phase-separated foci. The phase contrast channel (blue) and the mCherry channel (green) are merged. White arrows demarcate the same cellular location of the same focus over time. Blank arrows demarcate the cellular position now absent of a focus. Images are representative of four biological replicates. Scale bars: 2 µm. **b.** Cell elongation dissolves phase-separated foci. As in (a), except cells were treated with 10 µg/ml cephalexin to block cell division prior to spotting on agar pads.

**Figure S8.**
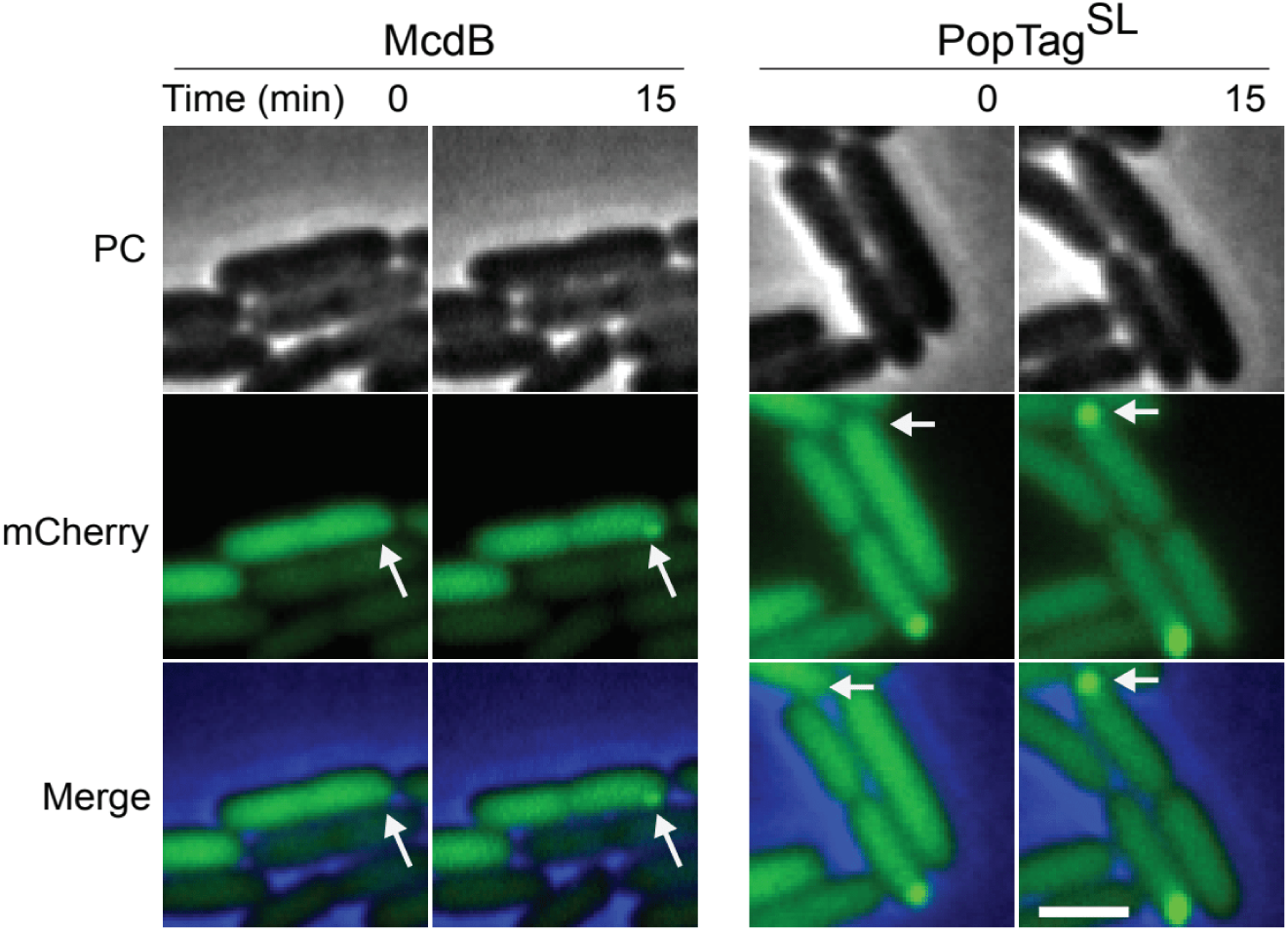
Focus formation after cell division. Phase contrast (PC) (gray), mCherry channel (green), and merged images are shown. White arrows demarcate the same cellular location before and after cell division. Images are representative of four biological replicates. Scale bar: 1 µm.

**Figure S9.**
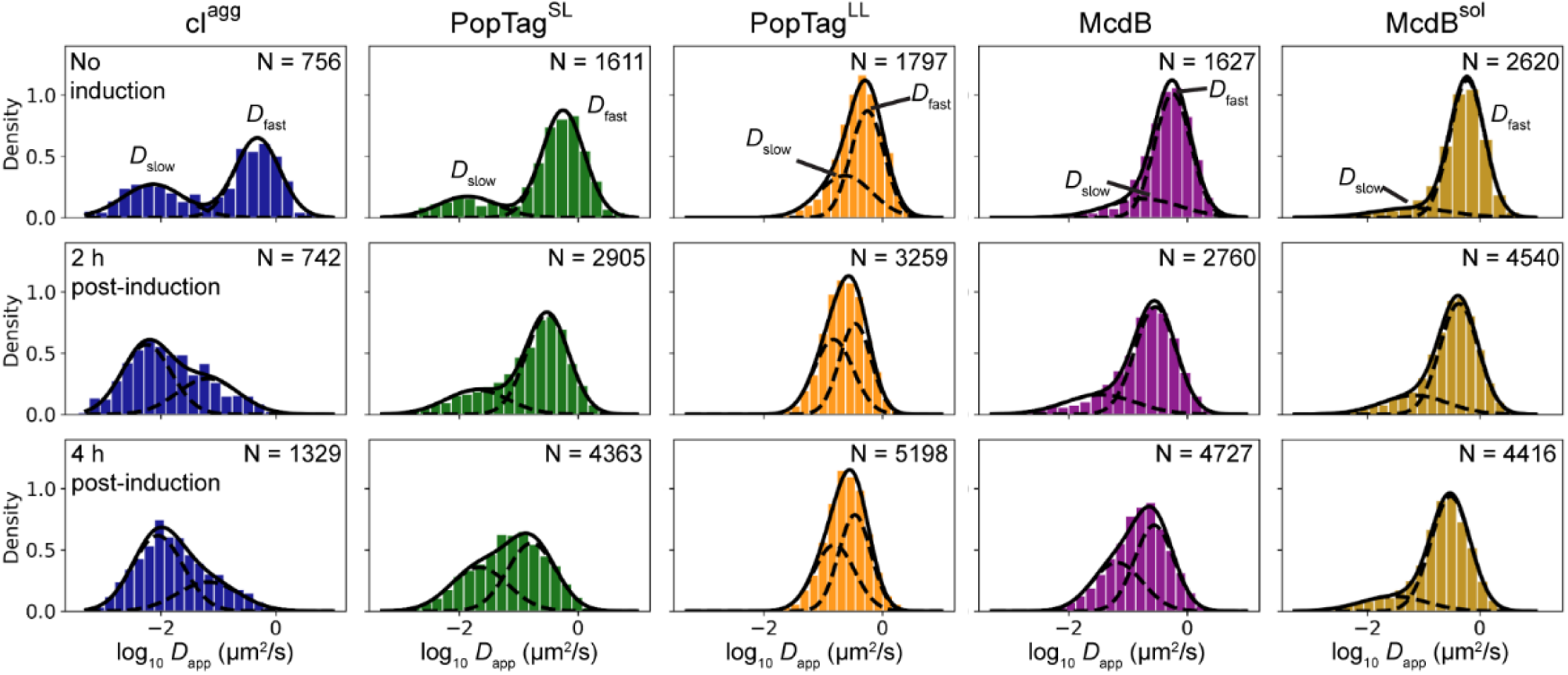
Diffusion coefficients of PAmCherry fusion proteins. Normalized log apparent diffusion coefficient histograms and two-component Gaussian mixture fit. The solid black line corresponds to the two-component Gaussian fit. The dashed black lines represent the Gaussian fit of each component which we refer to as *D*_app, slow_ and *D*_app, fast_.

**Figure S10.**
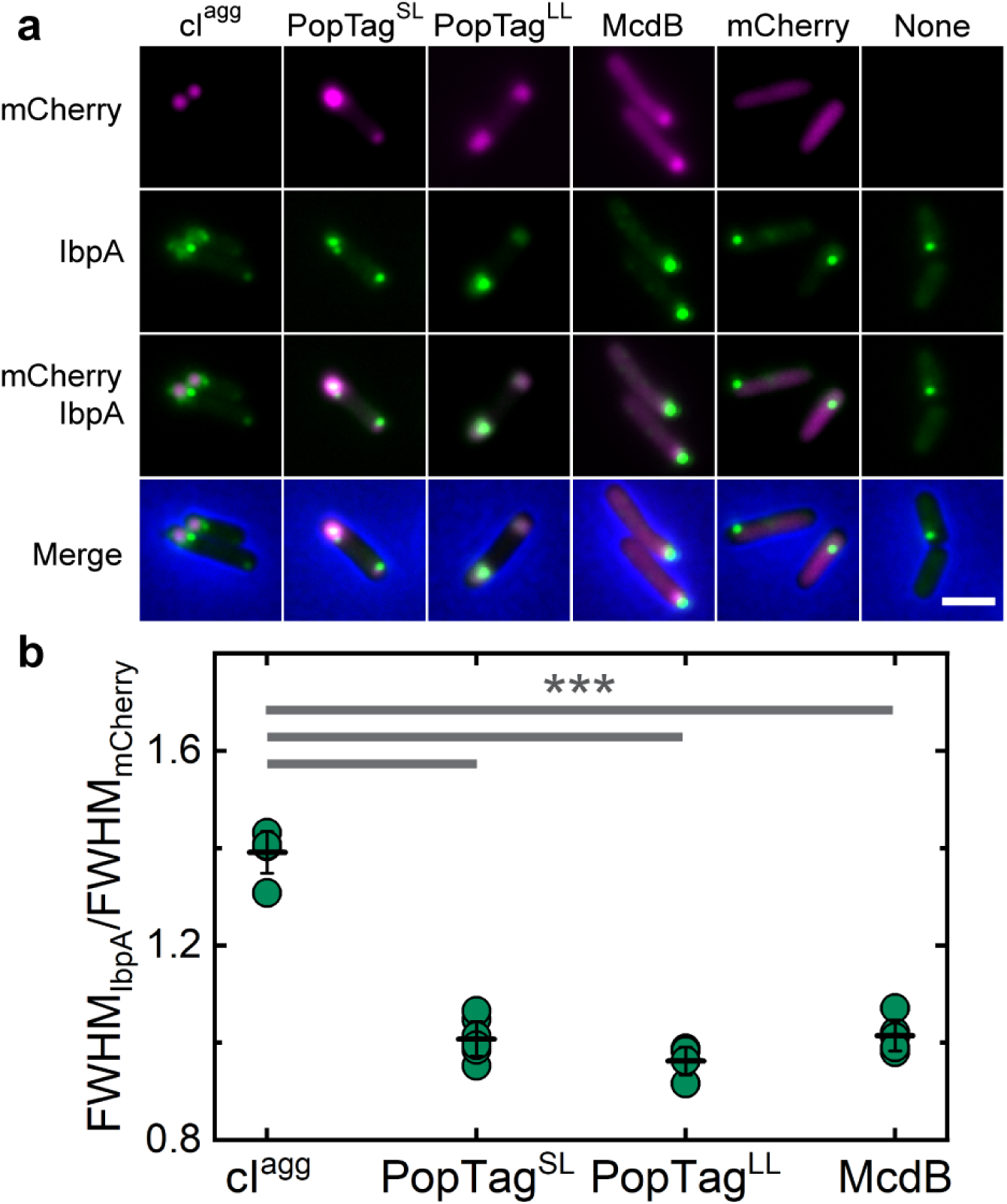
The nature of IbpA chaperone association with foci informs their material state *in vivo*. Wide-field fluorescence images of mCherry fusion foci (magenta) and IbpA-msfGFP foci (green). Phase Contrast is blue in the merge. Images are representative of three biological replicates. Scale bar: 2 µm. **b.** Quantification of the association of the foci of mCherry fusions and IbpA-msfGFP. Data points correspond to the ratio of the mean full-width at half maximum (FWHM) for the two channels (FWHM_IbpA_/FWHM_mCherry_) for the sum projections of technical replicates over three independent experiments. Black lines represent the mean ratio and the error bars represent the standard deviation. \*\*\**p* < 0.001 by student *t* test.

**Supplementary Table 1.** Strain and plasmids used in this study.

| Strains & Plasmids | Description | Source |
| --- | --- | --- |
| **Strains** |  |  |
| BL21 AE |  |  |
| BL21 AI |  |  |
| MG1655 $\Delta lacY$ | | |
| MG1655 $\Delta lacY$ *ibpA-msfGfp* | MG1655 encoding a C-terminal fusion of IbpA to msfGFP | This study |
| **Plasmids** |  |  |
|  | pET-*mNeonGreen-mcdB* | This study |
|  | pET-*His<sub>6</sub>-mNeonGreen-mcdB* | This study |
| pYH79 | pET-*His<sub>6</sub>-mCherry-mcdB* | This study |
| pTrc99A-*mCherry-cl78<sup>EP8</sup>* | pTrc99A expression vector allowing the inducible expression ($P_{trc}$ promoter) of a N-terminal fusion of cl78<sup>EP8</sup> (a mutagenized fragment of the lambda prophage repressor protein) to the mCherry fluorescent protein | <sup>28</sup> |
| pCA3 | pTrc99A-PAmCherry- *cl78<sup>EP8</sup>* | This study |
| pYH70 | pTrc99A-PAmCherry-*mcdB* | This study |
| pYH71 | pTrc99A-*mCherry-mcdB* | This study |
| pYH74 | pTrc99A-*mNeonGreen-mcdB* | This study |
| pYH75 | P<sub>trc</sub>99A-*mCherry-L6-PopTag* | This study |
| pYH76 | P<sub>trc</sub>99A-PAmCherry-L6-PopTag | This study |
| pYH77 | P<sub>trc</sub>99A-*mCherry* | This study |
| pYH78 | P<sub>trc</sub>99A-PAmCherry | This study |
| pYH80 | P<sub>trc</sub>99A-*mCherry-L78-PopTag* | This study |
| pYH88 | trc99A-PAmCherry-L78-PopTag | This study |
| pYH86 | P<sub>trc</sub>-*mCherry-mcdB<sup>sol</sup>* | This study |
| pYH88 | P<sub>trc</sub>-PAmCherry-*mcdB<sup>sol</sup>* | This study |

**Supplementary Table 2.**
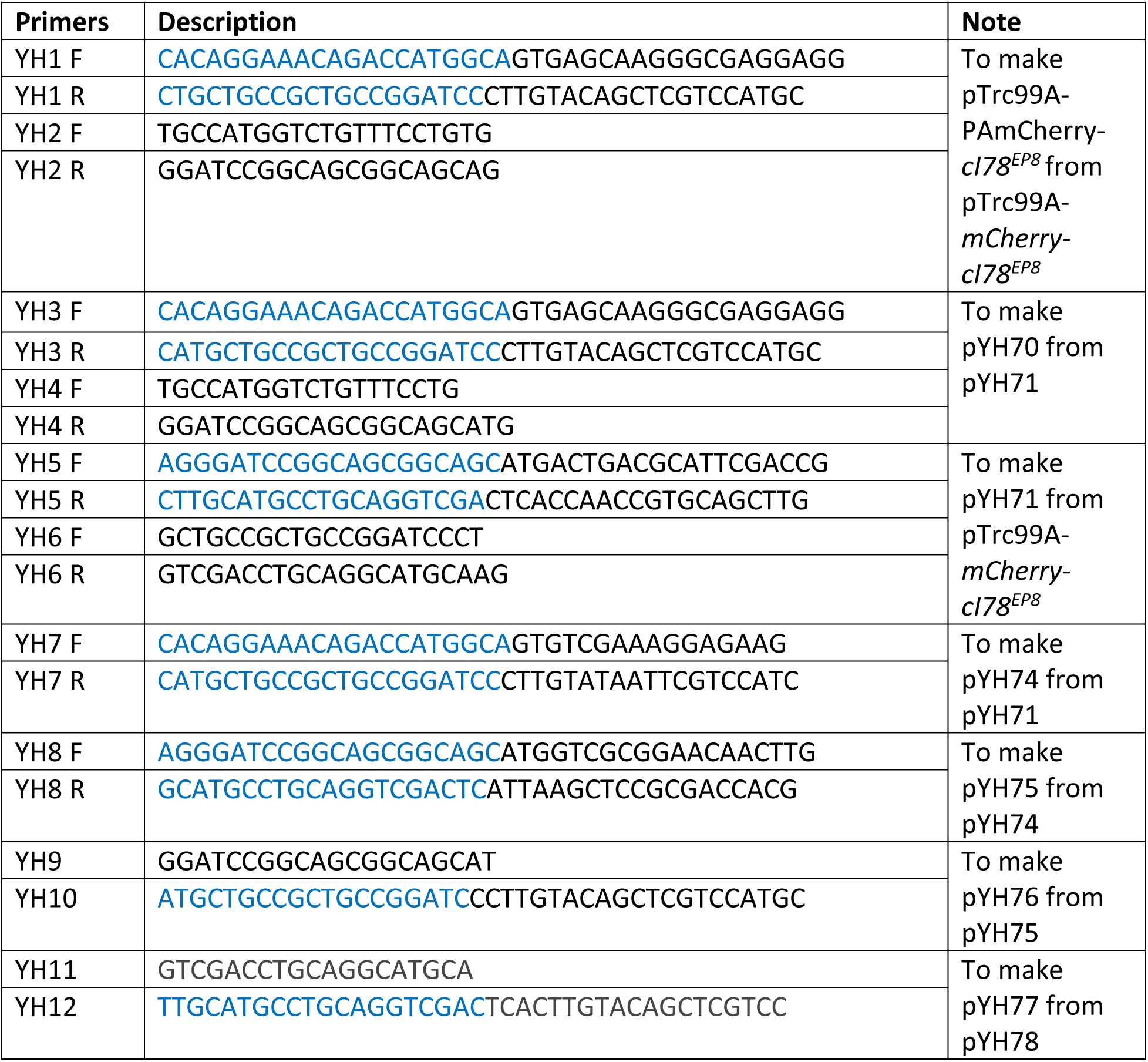
Primers used in this study.

**Supplementary Table 3.** Reagents used in this study.

| Reagent | Vendor | Catalog # | Use |
| --- | --- | --- | --- |
| 4-12% Bis-Tris NuPAGE gel | Thermo Fisher Scientific | NP0323BOX | SDS-PAGE |
| Bradford assay kit | Bio-Rad Laboratories | 5000006 | Measure protein content |
| Carbenicillin | Thermo Fisher Scientific | MT46100RG | Select for the plasmid in culture |
| DAPI | Thermo Fisher Scientific | D1306 | Label the nucleoid |
| Glucose | Thermo Fisher Scientific | 15023021 | Carbon source and inhibit protein expression from the $P_{trc}$ promoter |
| Glycerol | Thermo Fisher Scientific | G334 | Carbon source for *E. coli* MG1665 |
| Goat anti-rabbit IgG Secondary Antibody IRDye 800 CW | LI-COR | 926-32210 | Western Blot |
| IPTG | Thermo Fisher Scientific | 501126935 | Induce protein expression |
| L-arabinose | Thermo Fisher Scientific | AC365181000 | Induce protein expression |
| Mini-size polyvinylidene difluoride membrane | Bio-Rad | 1704156 | Protein transfer during Western Blot |
| Cephalexin | Thermo Fisher Scientific | AAJ6317206 | Inhibit cell division |
| A22 | Milipore sigma | SML0471 | The MreB inhibitor, which converts rod-shaped into round cells |
| Casamino acids | Milipore sigma | 65072-00-6 | AB medium |
| Thiamine | Milipore sigma | T4625-5G | AB medium |
| Uracil | Milipore sigma | 66-22-8 | AB medium |
| UltraPure agarose | Invitrogen | 16500100 | Agar pads |
| Ciprofloxacin | Thermo Fisher Scientific | AAJ6131714 | Compact the nucleoid |

**Supplementary Table 4.** Apparent diffusion coefficients and weight fraction resulting from two-state Gaussian mixture fitting.

| Protein | Sample induction (100 μM) | $D_{\text{app, slow}}$ ($\mu\text{m}^2/\text{s}$) | $\pi_{\text{slow}}$ | $D_{\text{app, fast}}$ ($\mu\text{m}^2/\text{s}$) | $\pi_{\text{fast}}$ |
| --- | --- | --- | --- | --- | --- |
| cl<sup>agg</sup> | None | $0.007 \pm 0.001$ | $0.38 \pm 0.02$ | $0.467 \pm 0.024$ | $0.62 \pm 0.02$ |
| | 2 h | $0.006 \pm 0.001$ | $0.62 \pm 0.03$ | $0.062 \pm 0.006$ | $0.38 \pm 0.03$ |
| | 4 h | $0.008 \pm 0.001$ | $0.65 \pm 0.03$ | $0.074 \pm 0.006$ | $0.35 \pm 0.03$ |
| PopTag<sup>SL</sup> | None | $0.013 \pm 0.001$ | $0.20 \pm 0.01$ | $0.572 \pm 0.016$ | $0.80 \pm 0.01$ |
| | 2 h | $0.024 \pm 0.002$ | $0.28 \pm 0.02$ | $0.314 \pm 0.010$ | $0.72 \pm 0.02$ |
| | 4 h | $0.023 \pm 0.001$ | $0.43 \pm 0.02$ | $0.165 \pm 0.006$ | $0.57 \pm 0.02$ |
| PopTag<sup>LL</sup> | None | $0.234 \pm 0.021$ | $0.37 \pm 0.04$ | $0.559 \pm 0.026$ | $0.63 \pm 0.04$ |
| | 2 h | $0.143 \pm 0.006$ | $0.47 \pm 0.03$ | $0.347 \pm 0.008$ | $0.53 \pm 0.03$ |
| | 4 h | $0.158 \pm 0.004$ | $0.46 \pm 0.03$ | $0.347 \pm 0.012$ | $0.54 \pm 0.03$ |
| McdB | None | $0.153 \pm 0.026$ | $0.23 \pm 0.05$ | $0.576 \pm 0.022$ | $0.77 \pm 0.05$ |
| | 2 h | $0.038 \pm 0.005$ | $0.23 \pm 0.02$ | $0.293 \pm 0.006$ | $0.77 \pm 0.02$ |
| | 4 h | $0.065 \pm 0.003$ | $0.40 \pm 0.02$ | $0.268 \pm 0.008$ | $0.60 \pm 0.02$ |
| McdB<sup>sol</sup> | None | $0.063 \pm 0.017$ | $0.14 \pm 0.01$ | $0.581 \pm 0.010$ | $0.87 \pm 0.01$ |
| | 2 h | $0.068 \pm 0.008$ | $0.23 \pm 0.01$ | $0.437 \pm 0.009$ | $0.77 \pm 0.01$ |
| | 4 h | $0.030 \pm 0.003$ | $0.19 \pm 0.01$ | $0.297 \pm 0.003$ | $0.81 \pm 0.01$ |

**Supplementary Table 5.**
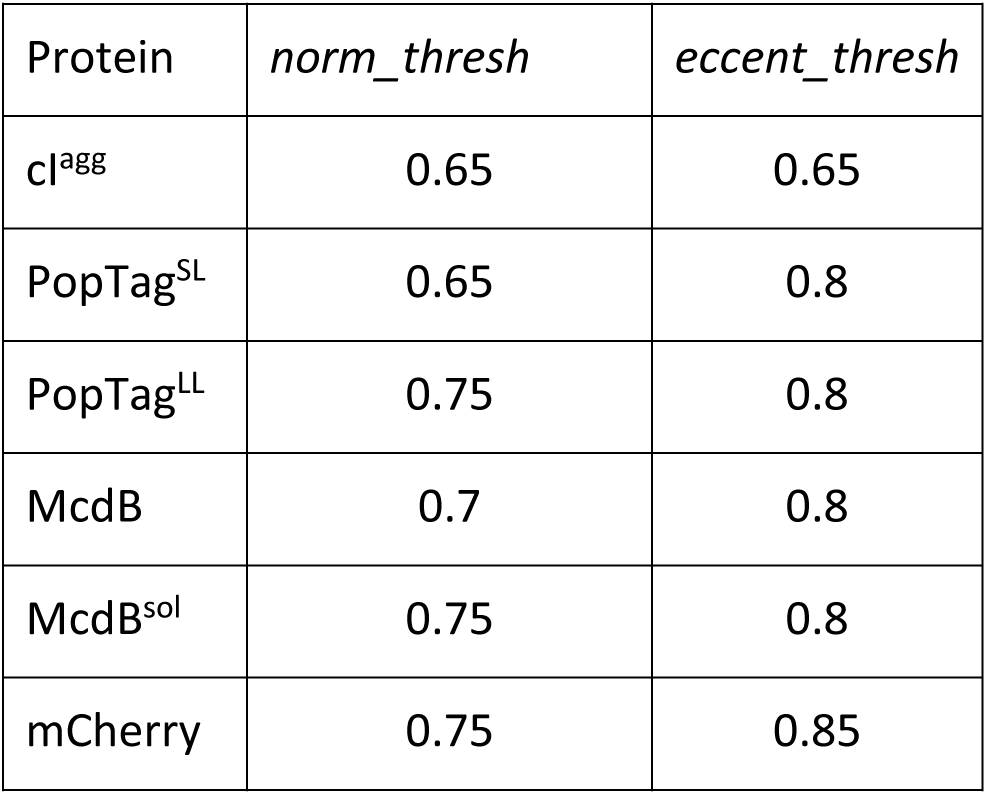
Parameters used for classification of cells in Condensation analysis.

| Protein | *norm_thresh* | *eccent_thresh* |
| --- | --- | --- |
| cI<sup>agg</sup> | 0.65 | 0.65 |
| PopTag<sup>SL</sup> | 0.65 | 0.8 |
| PopTag<sup>LL</sup> | 0.75 | 0.8 |
| McdB | 0.7 | 0.8 |
| McdB<sup>sol</sup> | 0.75 | 0.8 |
| mCherry | 0.75 | 0.85 |

**Video 1. Overexpressed mNG-McdB forms foci.** Indicated *E. coli* BL21 strains were induced with 1 mM IPTG for expression of mNG-McdB. Time-lapse movies from 0 to 4 h post-induction were recorded. The phase contrast channel is shown in gray and the mNG channel is in green. This video is representative of three biological replicates. Scale bar: 5 µm.

**Video 2. Effects of temperature shift on focus stability.** Cells that formed foci at 25°C were transitioned to 37°C in a stage-top incubator. Scale bar: 1 µm

**Video 3. Effects of changing cell volume on focus stability.** *E. coli* cell volume was increased by treating cells with the MreB inhibitor, A22 (10 µg/ml). Scale bar: 2 µm.

**Video 4. Localized lysis of the cell.** One cell pole was lysed using a high intensity laser. The time-lapse video shows solubilization of the opposing focus and shift of fluorescent signal to the opposite end. Scale bar: 1 µm.

**Video 5. mCherry-McdB^sol^ condenses by repulsive interactions with the nucleoid.** mCherry-McdB (left) and mCherry-McdB^sol^ (right) foci in *E. coli* at 4 h post-induction. Cells were treated with ciprofloxacin (50 mM) for nucleoid compaction and stained with DAPI (2 µM) for nucleoid visualization (bottom). Merged images of mCherry channel (green), DAPI (magenta), and Phase contrast (blue) are shown. The video is representative of three biological replicates. Scale bars: 1 µm.

**Video 6. Generational-dilution dissolves condensates.** Phase contrast (gray), mCherry channel (green), and merged images are shown. Images are representative of four biological replicates. Scale bar: 2 µm.

**Video 7. Cell elongation dissolves phase-separated condensates.** Cells were treated with 10 µg/ml cephalexin to block cell division. Phase contrast (gray), mCherry channel (green), and merged images are shown. Images are representative of four biological replicates. Scale bar: 2 µm.

